# Heterologous production of 1-tuberculosinyladenosine in *Mycobacterium kansasii* models pathoevolution towards the transcellular lifestyle of *Mycobacterium tuberculosis*

**DOI:** 10.1101/2020.04.25.060590

**Authors:** Marwan Ghanem, Jean-Yves Dubé, Joyce Wang, Fiona McIntosh, Daniel Houle, Pilar Domenech, Michael B. Reed, Sahadevan Raman, Jeffrey Buter, Adriaan J. Minnaard, D. Branch Moody, Marcel A. Behr

## Abstract

*Mycobacterium kansasii* is an environmental non-tuberculous mycobacterium that causes opportunistic tuberculosis-like disease. It is one of the most closely related species to the *M. tuberculosis* complex. Using *M. kansasii* as a proxy for the *M. kansasii-M. tuberculosis*-common ancestor, we asked whether introducing the *M. tuberculosis*-specific gene pair *Rv3377c-Rv3378c* into *M. kansasii* affects the course of experimental infection. Expression of these genes resulted in the production of an adenosine-linked lipid species, known as 1-tuberculosinyladenosine (1-TbAd), but did not alter growth *in vitro* under standard conditions. Production of 1-TbAd enhanced growth of *M. kansasii* under acidic conditions through a bacterial cell-intrinsic mechanism independent of controlling pH in the bulk extracellular and intracellular spaces. Production of 1-TbAd led to greater burden of *M. kansasii* in the lung of C57Bl/6 mice during the first 24 hours after infection and *ex vivo* infections of alveolar macrophages recapitulated this phenotype within the same time frame. However, in long-term infections, production of 1-TbAd resulted in impaired bacterial survival in both C57Bl/6 mice and *Ccr2-/-* mice. We have demonstrated that *M. kansasii* is a valid surrogate of *M. tuberculosis* to study virulence factors acquired by the latter organism, yet shown the challenge inherent to studying the complex evolution of mycobacterial pathogenicity with isolated gene complementation.

**IMPORTANCE:** This work sheds light on the role of the lipid 1-tuberculosinyladenosine in the evolution of an environmental ancestor to *M. tuberculosis*. On a larger scale, it reinforces the importance of horizontal gene transfer in bacterial evolution and examines novel models and methods to provide a better understanding of the subtle effects of individual *M. tuberculosis*-specific virulence factors in infection settings that are relevant to the pathogen.

## INTRODUCTION

*M. tuberculosis* virulence factors have been established by genetic knock-out and complementation within the pathogen, producing evidence of an attenuation of virulence in *ex vivo* or *in vivo* experimental infections (1, 2). While numerous virulence-associated loci have been identified with this approach, the majority of these are intact in the genomes of non-transmissible environmental mycobacteria, such as *M. kansasii* (3–6). *M. kansasii* is readily isolated in clinical settings from pulmonary infections and we have previously shown that it can be studied in an experimental lung model (3). However, although it causes TB-like disease, *M. kansasii* infections disproportionately affect patients with underlying pulmonary diseases or immunosuppression, and there is no evidence supporting its transmission between individuals (7, 8). The conservation of many virulence factors across *M. tuberculosis* and *M. kansasii*, including the ESX-1 secretion system, PhoPR 2-component system and DosR/S/T regulon suggest that they play a role in a broader survival strategy used by mycobacteria (9–11). For example, some of these factors may be needed for survival of *M. kansasii* within free-living phagocytic amoeba, but their presence does not provide *M. kansasii* with the pathogenic capabilities of *M. tuberculosis* inside human hosts (12). Consequently, there is currently an incomplete understanding of how *M. tuberculosis* emerged as a human-adapted professional pathogen.

There has been a growing body of evidence over the past decade showing that HGT events have happened during mycobacterial speciation and are associated with the step-wise emergence of pathogenic species (13–16). Fifty-five genes have been acquired by *M. tuberculosis* since its divergence from the *M. kansasii-M. tuberculosis* common ancestor (MKMTCA) (13). Although many of these HGT genes have no postulated function, the *Rv3376-Rv3378c* genomic island uniquely present in *M. tuberculosis* is known to encode a class II terpene cyclase (*Rv3377c*) and a tuberculosinyl transferase (*Rv3378c*). Together, the two enzymes are responsible for the conversion of geranylgeranyl pyrophosphate (GGPP) into the recently identified *M. tuberculosis*-specific lipid 1-tuberculosinyladenosine (1-TbAd), which is a potential diagnostic molecular marker for TB disease (17–20). 1-TbAd further undergoes a chemical rearrangement, known as the Dimroth reaction, to generate *N*^6^-TbAd (18). While GGPP is found in both species and used as an intermediate in the biosynthesis of 1-TbAd by *M. tuberculosis* (17, 19), it is part of the biosynthetic pathway for the production of carotenoid pigments of *M. kansasii*, giving its characteristic yellow colour (21).

We previously reported the important role of 1-TbAd in protecting *M. tuberculosis* from phagosomal acidification inside macrophages (22). In the present work, we have characterized the effect of 1-TbAd production in *M. kansasii* complemented with *Rv3377c and Rv3378c*. Here, we show that *in vitro* growth kinetics and colony morphology in liquid and on solid media are unaltered by *Rv3377c-Rv3378c* expression. 1-TbAd confers a growth advantage in acidic media compared to wild-type *M. kansasii*, which we further demonstrated to be independent of cytosolic and culture medium pH control, suggesting a compartmental mechanism of protection for the bacterium. *Rv3377c-Rv3378c* provided an early advantage to bacterial replication during pulmonary infection in mice, consistent with enhanced survival in alveolar macrophages. However, the *M. kansasii:Rv3377-78c* was outcompeted by the wild-type during long-term murine infection. This study demonstrates that we can use *M. kansasii* as a proxy of the MKMTCA to explore the complex evolution of *M. tuberculosis*. It also shows that gene acquisition likely provided advantages for specific contexts, including a possible and unexpected role in survival within alveolar macrophages during early stages of infection, despite tradeoffs or challenges under other circumstances requiring further evolution to overcome.

## RESULTS

### Expression of *Rv3377c-Rv3378c* in *M. kansasii* leads to 1-TbAd production

We introduced the *M. tuberculosis*-specific gene pair *Rv3377c-Rv3378c* into the *M. kansasii* genome within an integrative plasmid containing hygromycin resistance to produce *M. kansasii::Rv3377-78c* (see methods). As a control for subsequent experiments, an integrative ‘empty vector’ (EV) was employed. After labelling with ^14^C-adenosine and lipid extraction using chloroform and methanol, radio-thin-layer chromatography (TLC) was used to detect adenosine-linked lipids in *M. kansasii:Rv3377-78c* clones for comparison to *M. tuberculosis* strain H37Rv (Fig. 1a). Conventional molybdenum-based sprays followed by charring broadly detected all lipids as a loading control, suggesting a lack of broad lipid changes detectable at the TLC level after gene transfer (Fig. 1b). Whereas uncomplemented bacterial extracts showed material at the origin and one weak band in radio-TLC, *Rv3377c-Rv3378c* complementation generated at least 5 additional lipid species. Three of these novel lipids co-migrated with compounds from *M. tuberculosis* strain H37Rv. Both results strongly suggested the successful genetic transfer of *M. tuberculosis*-associated adenosine-linked lipids to *M. kansasii*. 5% phosphomolybdic acid reagent (PMA) staining showed that similar amounts of total lipids were spotted for each *M. kansasii* sample (Fig. 1b).

**Figure 1 –.**
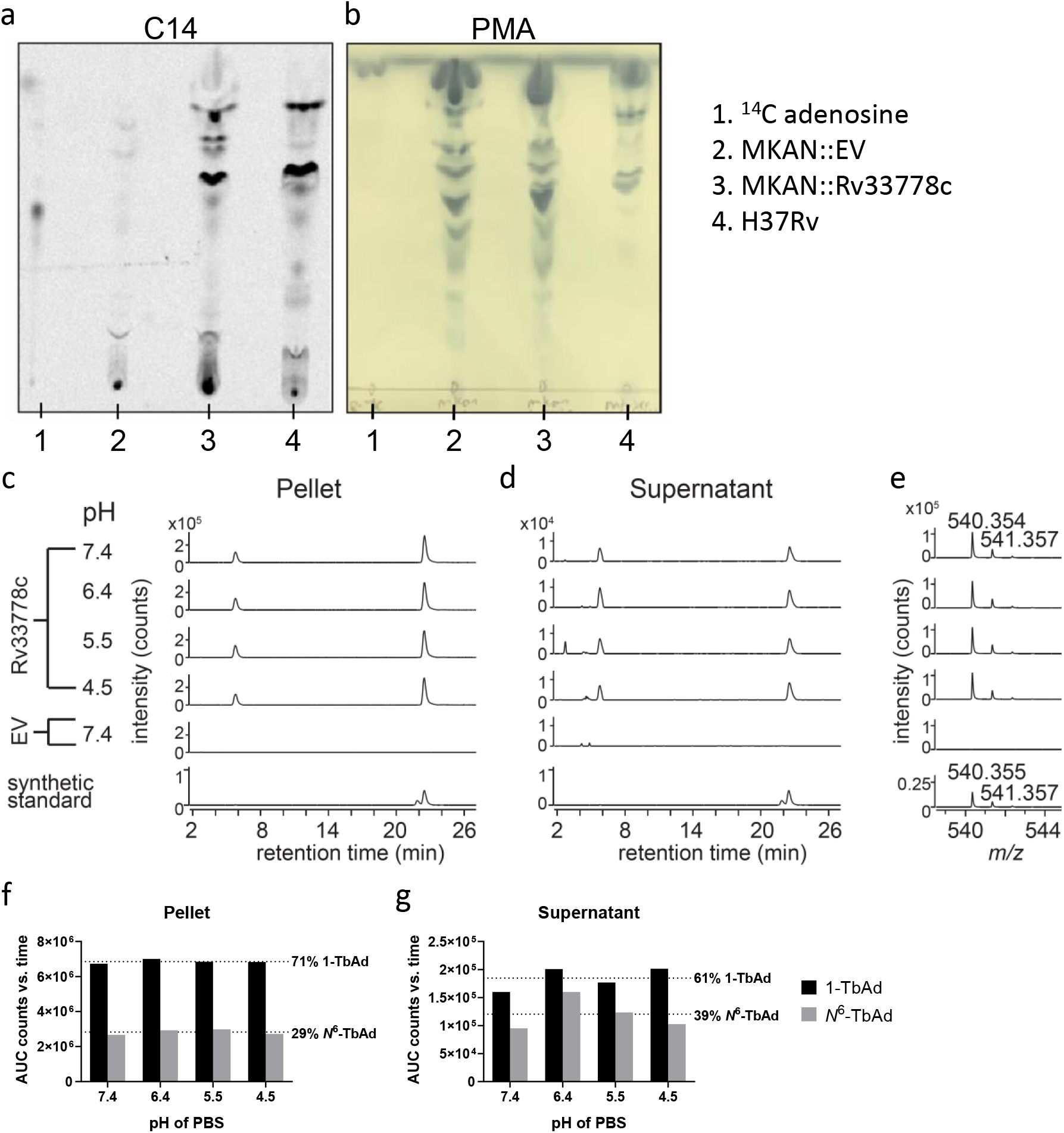
*M. kansasii::Rv3377-78c* produces adenosine-linked lipids 1-TbAd and N6-TbAd. (a) Detection of adenosine-linked lipids extracted from *M. kansasii*::EV (MKAN::EV), *M. kansasii::Rv3377-78c* (MKAN::Rv33778c) and *M. tuberculosis* (H37Rv) through radiolabelling and separation using normal-phase silica thin-layer chromatography. (b) Visualization of migration pattern of total lipids from each sample after staining with 5% phosphomolybdic acid reagent. (c-e) Lipids from *M. kansasii* derived from cell pellets or culture supernatant incubated for two hours at the indicated pH, neutralized and then extracted with organic solvent. Product was analyzed in comparison with a synthetic standard for 1-TbAd, where the slightly later and larger peak corresponds to native 1-TbAd from *Mtb*. (e) The mass spectrum of lipids extracted from 21-22 min for *M. kansasii* show a m/z value that matches with the measured and expected mass of a 1-TbAd standard (21) (f-g) Total extracted lipids expressed as area under curve from counts versus retention time of the extracted ion chromatogram. 1.0 μM of synthetic 1-TbAd was used as the standard.

High-performance liquid chromatography-mass spectrometry (HPLC-MS) was used to chemically identify the compounds produced by *M. kansasii::Rv3377-78c*, in which the expected retention of 1-TbAd (22.7 min) and *N*^6^-TbAd (5.8 min) were known (Fig. 1c-g). Whereas the *M. kansasii*::EV control did not release compounds that comigrated with either 1-TbAd or *N*^6^-TbAd, *M. kansasii:Rv3377-78c* produced high intensity (6.7-7.0 x10^6^ counts) signals (m/z 540.354) matching the expected retention time and mass (m/z 540.3544) of the proton adducts of 1-TbAd. The extractions were performed at a range of pH (4.5 – 7.4) since both the Dimroth reaction that generates *N*^6^-TbAd and the capture of lysosomotropic agents are sensitive to pH, as previously explained (18, 22, 23). Similar to results with *M. tuberculosis* in which 9 % of the TbAd pool was released (22), we observed stronger (~ 10-fold) signals in the pellet compared to the supernatant. However, there was no clear impact of altering pre-extraction pH for two hours to the release nor relative abundance of 1-TbAd and *N*^6^-TbAd, and thus such effects, if existent, did not occur under the tested conditions (Fig. 1c-g). Similar to patterns observed from *M. tuberculosis*, more 1-TbAd than *N*^6^-TbAd was recovered from *M. kansasii::Rv3377-78c* (Fig. 1f-g). In 4 tested cultures of *M. kansasii::Rv3377-78c*, 1-TbAd represented 0.125 +/- 0.04% of the total lipid mass (vs. 0.76% in *M. tuberculosis*) (data not shown).

### 1-TbAd production does not change visible characteristics of *M. kansasii* in conventional media

To characterize any overt phenotypic effects of 1-TbAd production on *M. kansasii, we* assessed its influence on *in vitro* characteristics of the bacterial culture. *M. kansasii::Rv3377-78c* grew similarly to wild-type *M. kansasii* and *M. kansasii*::EV in 7H9 (Fig. 2a) and on 7H10 agar (Fig. 2b). Carotenoid pigments get integrated into bacterial cell membranes, maintain membrane fluidity and provide support against external stressors (24). Since the production of 1-TbAd requires the same intermediate GGPP as that of the yellow pigment that *M. kansasii* produces when exposed to light, we then tested *M. kansasii::Rv3377-78c*’s ability to turn yellow after light exposure in order to rule out the possibility that a pigment-related phenomenon might affect our outcomes (Fig. 2b). *M. kansasii::Rv3377-78c* retained photochromogenic abilities by turning yellow after exposure to light at room temperature within the same timeframe as *M. kansasii*::EV.

**Figure 2 –.**
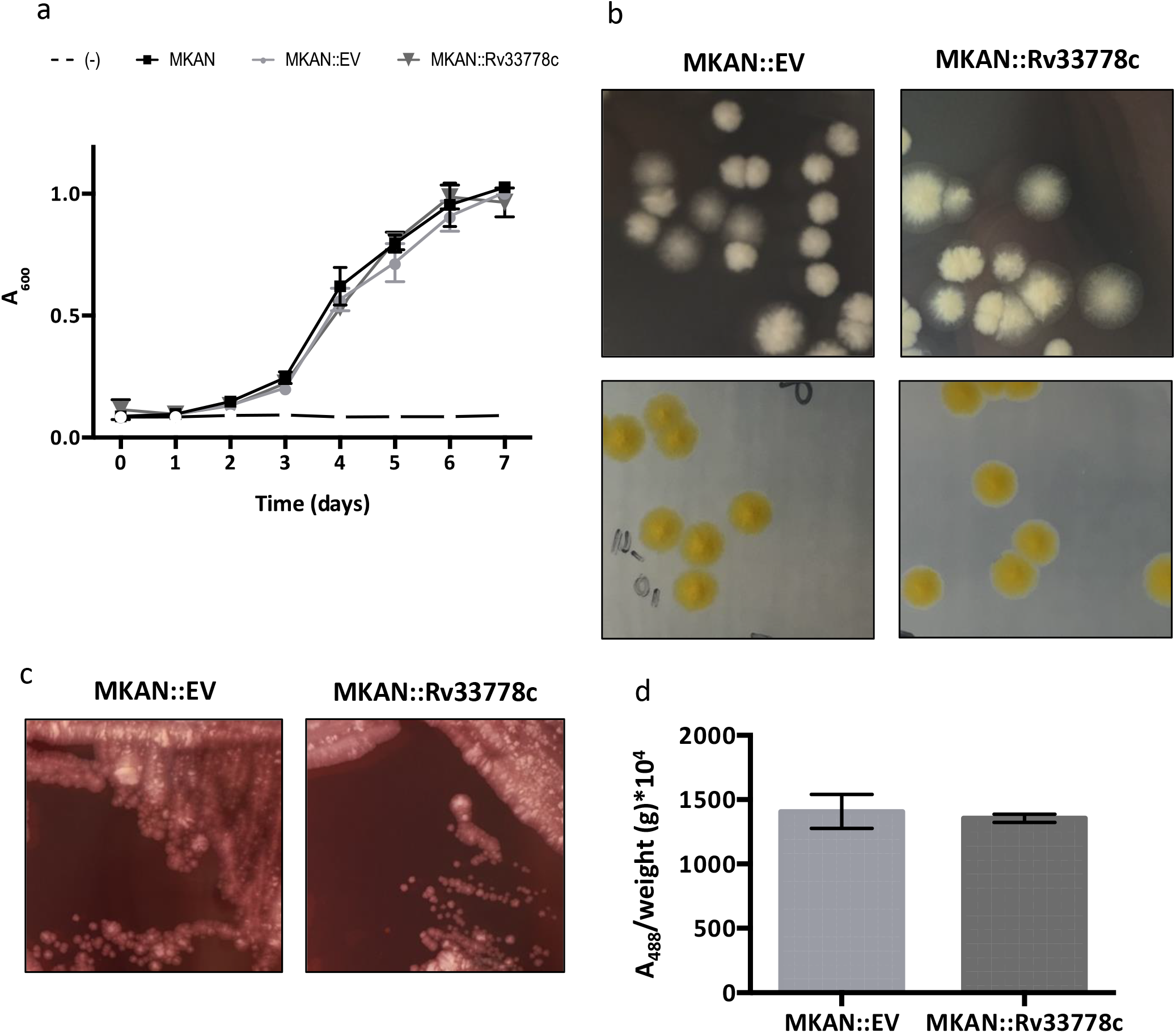
1-TbAd production does not influence the *in vitro* growth characteristics and behaviours of *M. kansasii*. (a) Comparative OD_600_ growth kinetics of wild-type *M. kansasii* (MKAN), *M. kansasii*::EV (MKAN::EV) and *M. kansasii::Rv3377-78c* (MKAN::Rv33778c) at 37°C in 7H9 broth. The data are presented as the mean of technical triplicates ± SD. The data are representative of three independent experiments. (b) Colony morphology in 2 different incubation settings of *M. kansasii*::EV (MKAN::EV) and *M. kansasii::Rv3377-78c* (MKAN::Rv33778c) on 7H10 plates. (c) Colony morphology of *M. kansasii*::EV (MKAN::EV) and *M. kansasii::Rv3377-78c* (MKAN::Rv33778c) on 7H10 plates supplemented with Congo Red. (d) Quantitative analysis of Congo Red dye retention by *M. kansasii*::EV (MKAN::EV) and *M. kansasii::Rv3377-78c* (MKAN::Rv33778c). DMSO extraction was followed by absorbance at 488nm divided by the weight of the dry culture pellet after washing (in grams). The data are plotted as the mean of technical triplicates ± SD. The data are representative of two independent experiments.

Congo Red is an amphiphilic dye that binds to the mycobacterial cell membrane. When grown on Congo Red-containing 7H10 plates, different mycobacteria retain the dye distinctly, and this feature has been associated with differences in the interactions of bacterial cells within the colony (25). In this study, *M. tuberculosis* absorbed the dye and became red, while *M. kansasii* colonies remained white on the plate. Since 1-TbAd is found on the cell surface and has an amphipathic character (17), we tested the effect of its production on intra-colony bacterial interactions of *M. kansasii::Rv3377-78c*. Visual inspection of colonies (Fig. 2c) and absorbance at 488nm (Fig. 2d) showed no difference in retention of Congo Red between *M. kansasii*::EV and *M. kansasii::Rv3377-78c*.

### 1-TbAd production enhances growth of *M. kansasii* in acidic media

The ability to survive in mildly acidic environments is a key feature of mycobacteria, both environmental and pathogenic (26, 27). It was recently shown that 1-TbAd production confers a growth advantage over a pH range (5.0-5.4) comparable to that of an activated phagolysosome, which is not tolerated by most bacteria (22). As 1-TbAd can be shed extracellularly to de-acidify the phagosomal environment as seen in *M. tuberculosis, we* hypothesized that the production of 1-TbAd by *M. kansasii::Rv3377-78c* may modulate media pH (22). As expected, *M. kansasii::Rv3377-78c* was able to grow in lower pH than *M. kansasii*::EV in 7H9 culture media (Sup. Fig. 1). However, at 8 and 17 days post inoculation, both *M. kansasii*::EV and *M.kansasii::Rv3377-78c* slightly increased the pH of the media where there was bacterial growth, and to a similar extent (Fig. 3). Therefore, although we observed enhanced growth with *M. kansasii::Rv3377-78c* compared to *M. kansasii*::EV, we could not demonstrate a causative role for 1-TbAd raising the extracellular pH of the culture media under these conditions.

**Figure 3 –.**
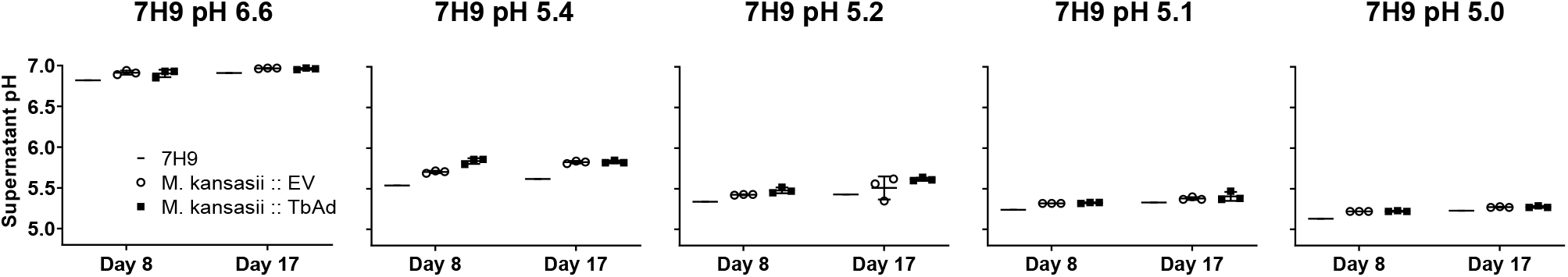
1-TbAd production enhances growth at low pH where growth is associated with culture medium alkalization. *M. kansasii*::EV (MKAN::EV) and *M. kansasii::Rv3377-78c* (MKAN::Rv33778c) cultures were inoculated at equal OD_600_ into fresh pH-adjusted 7H9 (using HCl titration) and incubated at 37°C in a rolling incubator over 17 days. The starting pH of the cultures is indicated above each graph. The pH of the supernatant was measured at days 8 and 17. The data are presented as the mean of technical triplicates ± SD. The data are representative of three independent experiments.

### Synthetic 1-TbAd does not directly enhance growth of *M. kansasii*

Prior work with *M. tuberculosis* estimated that 1-TbAd might naturally accumulate to μM concentrations in phagosomes, and 5-20 μM 1-TbAd alters lysosomal pH and morphology in human macrophages (22). The proposed lysosomotropic mechanism requires that 1-TbAd access a low pH compartment where the uncharged conjugate base binds protons to raise pH and regenerate a concentration gradient that promotes further entry of uncharged conjugate base to the acidic compartment. Whereas this mechanism can relieve pH stress on the bacterium, the major alternative, which is not exclusive of lysosomotropism, is that 1-TbAd directly signals for bacterial growth and division. To distinguish these mechanisms we ‘chemically complemented’ WT *M. kanasii* with synthetic 1-TbAd and *N*^6^-TbAd added externally in media. In this experiment TbAd (already carrying a proton) should not alter pH, but would contact bacteria in high concentrations. As expected the addition of synthetic 1-TbAd (p*K*_a_ ~ 8.5) and *N*^6^-TbAd (p*K*_a_ ~ 3.8) did not alter the pH of the 7H9 media (Sup. Fig. 2a). Next we inoculated *M. kansasii* into pH-adjusted 7H9 broth (pH 4.0, 4.8, 5.0, 5.2, 5.4, 6.7,) containing 1, 5, 10 or 20 μM TbAd and monitored growth over 16 days (Fig. 4 and Sup. Fig. 2b). With increasing doses of 1-TbAd or its isomer *N*^6^-TbAd, we did not observe any promotion of *M. kansasii* growth in normal nor acidic 7H9 broth (Fig. 4 and Sup. Fig 1b). In fact, 1-TbAd partially inhibited growth at 20 μM, the highest dose tested, at normal pH. These data demonstrated that the protection from low pH in 7H9 culture media afforded by *Rv3377c-Rv3378c* complementation in *M. kansasii* is cell-intrinsic, promotes growth only at low pH and does not occur with direct exposure to protonated TbAd.

**Figure 4 –.**
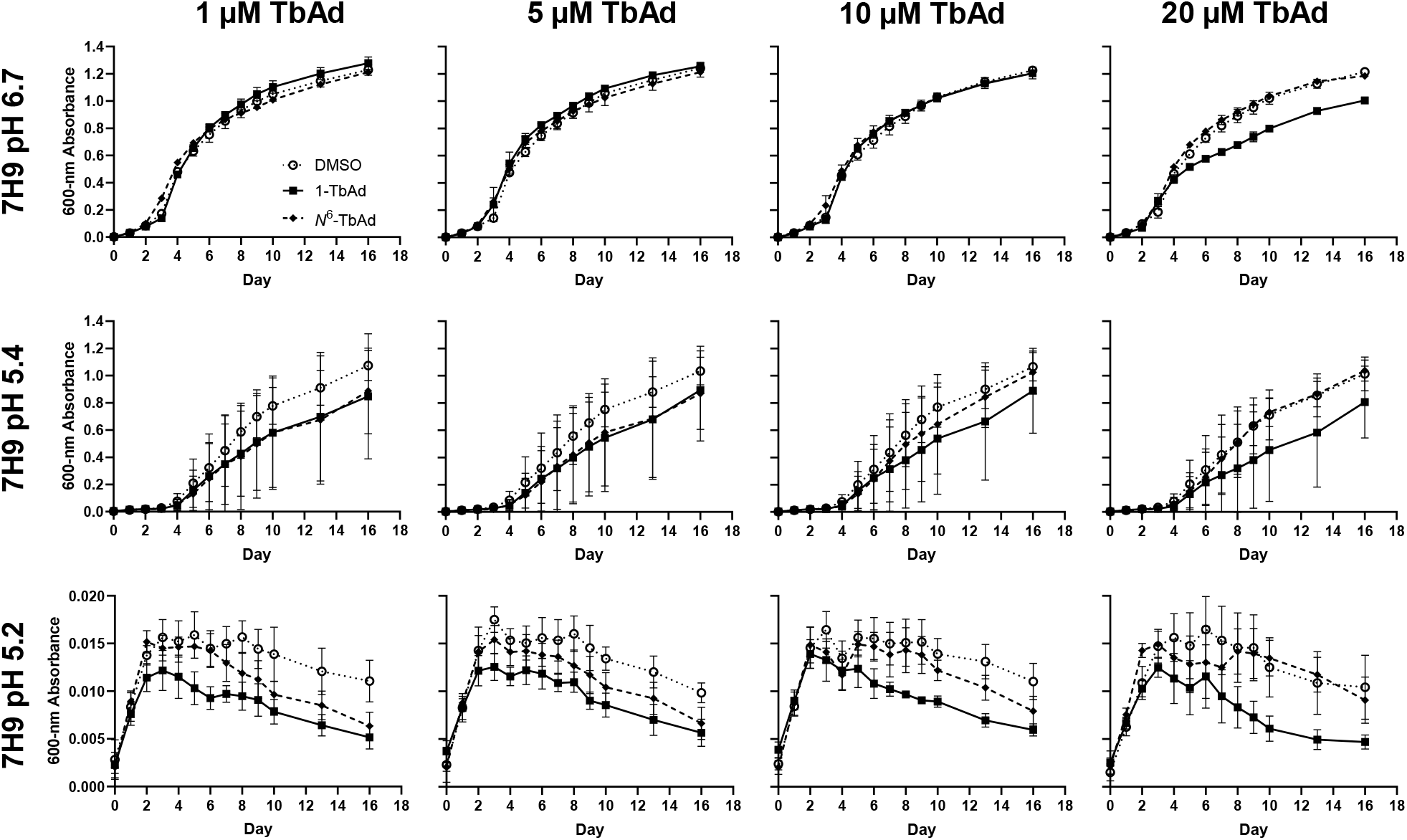
Chemical complementation of *M. kansasii* with synthetic TbAd does not promote growth at any pH. (a) *M. kansasii* WT was inoculated into fresh pH-adjusted 7H9 (using HCl titration) containing the indicated concentration of TbAd isomer or DMSO, and then incubated at 37°C in 96-well plates over 16 days. OD_600_ was measured every 1-3 days. The data are presented as the mean of technical quadruplicates ± SD.

### 1-TbAd does not maintain bulk cytosolic pH to enhance *M. kansasii* growth in low pH media

We aimed to identify whether 1-TbAd production alters/maintains the pH of the bacterial cytosol when grown in acidic media. *M. kansasii*::EV or *M.kansasii::Rv3377-78c* were stained with carboxyfluorescein diacetate succinimidyl ester (CFSE) to measure their intracellular pH while monitoring their growth in different pH-adjusted 7H9 broth (pH 4.0, 4.8, 5.0, 5.2, 5.4, 6.0, 6.7, 7.2) overnight (Fig. 5). Lower initial pH of the media was associated with lowering intracellular pH over the experimental timeframe, as expected. Importantly, we observed the expected growth advantage with 1-TbAd production at lower pH (5.0), but there was no apparent intracellular pH difference between *M. kansasii* strains at this or any pH. Thus, 1-TbAd production does not aid *M. kansasii* growth at low pH by maintaining the pH of the bulk intracellular milieu, suggesting the growth advantage is provided by countering the effect of low pH in a specific region of the cell.

**Figure 5 –.**
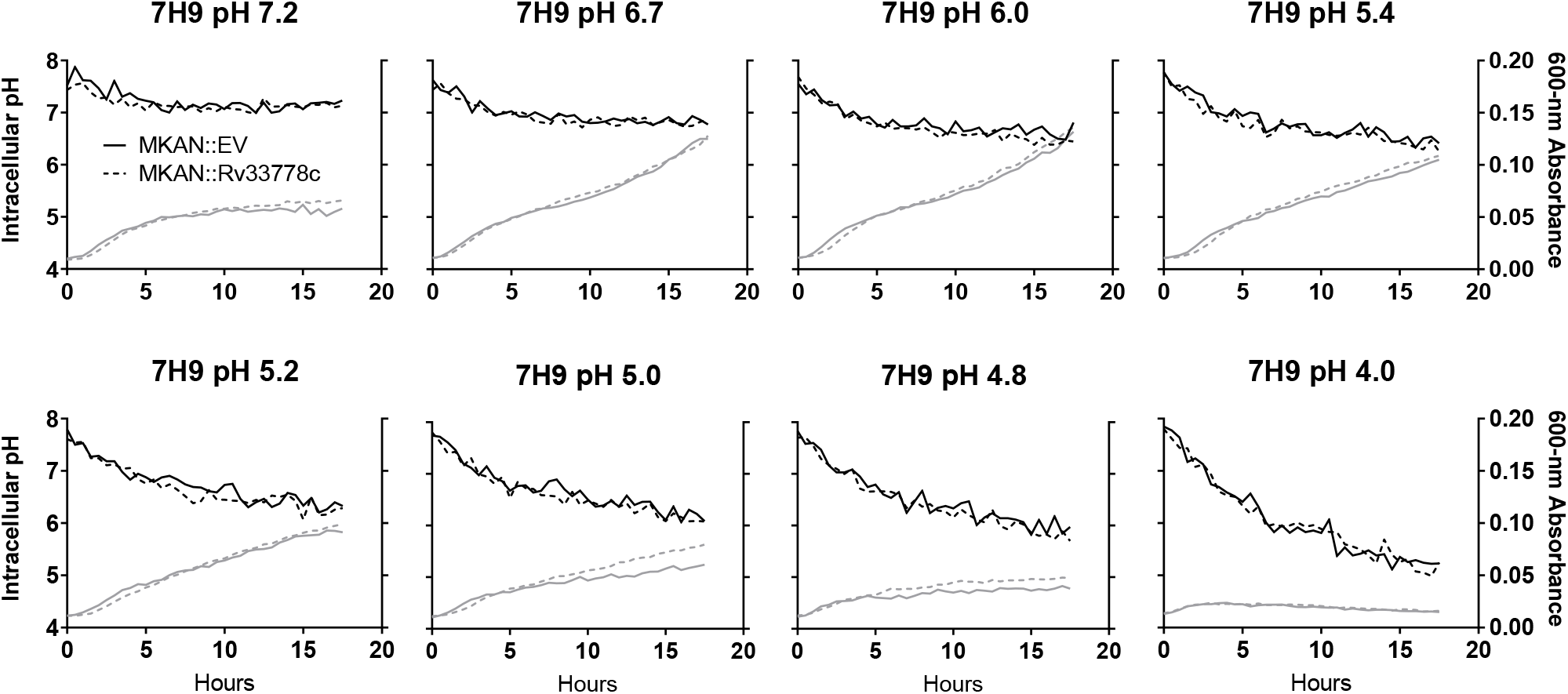
1-TbAd does not alter the intracellular pH of bacteria. *M. kansasii*::EV (MKAN::EV - solid lines) and *M. kansasii::Rv3377-78c* (MKAN::Rv33778c - dashed lines) cultures at equal OD_600_ were stained with CFSE, inoculated into fresh pH-adjusted 7H9 and incubated shaking at 37°C in 96-well plates placed in the dark. Growth (grey / OD_600_ measurements) and intracellular pH (black / pH calculated from fluorescence excitation-emission ratios) readings were taken at 30-minute intervals overnight. The data are presented as the median of technical triplicates. Data are representative of 5 independent experiments.

### 1-TbAd production enhances establishment of *M. kansasii* lung infection and survival in alveolar macrophages

1-TbAd is hypothesized to promote virulence of *M. tuberculosis* by countering phagosome acidification, enhancing survival of the pathogen *in cellulo* (19, 28, 29). We therefore wished to test the virulence of *M. kansasii::Rv3377-78c*. In C57Bl/6 mice, *M. tuberculosis* expands in a logarithmic scale within the early course of infection. In sharp contrast, *M. kansasii* remains at its initial levels of infection, suggesting it is a good model for acquisition-of-virulence studies within the mouse (3). As a measure of virulence, we hypothesized that the pulmonary bacterial load of *M. kansasii* in mice would be enhanced with 1-TbAd production. During pilot experiments, we infected mice with *M. kansasii*::EV and *M. kansasii::Rv3377-78c* through aerosolization, and infections with either strain resulted in a pulmonary burden within one log of the initial infection up to day 42, with no clear differences between both groups with the sample sizes used (Sup. Fig. 3).

Despite repeated attempts to standardize the inoculum, we consistently noted a higher number of 1-TbAd-producing *M. kansasii::Rv3377-78c* one day after aerosol infection when compared with *M. kansasii*::EV (not shown). To test if *Rv3377c-Rv3378c* was altering the dose administered or instead enhancing establishment or early growth, mice were aerosolized with *M. kansasii*::EV or *M. kansasii::Rv3377-78c* and their lungs were collected and homogenized shortly (4 hours) after infection and compared to bacterial counts 24 hours after infection (Fig. 6a-b). Equivalent 4-hour CFU counts were observed for *M. kansasii*::EV and *M. kansasii::Rv3377-78c*; only the latter multiplied successfully 24 hours later (Fig. 6a). *M. kansasii::Rv3377-78c* showed a 1.5-fold increase in numbers from 4 to 24 hours while *M. kansasii*::EV numbers remained equal (Fig. 6b). The experiment was performed three times, once at a low dose of 100 CFUs/lung and twice at a higher dose of 750 CFU/lung, with *M. kansasii::Rv3377-78c* consistently attaining larger numbers by 24 hours post-infection in all three instances (Sup. Fig. 4).

**Figure 6 –.**
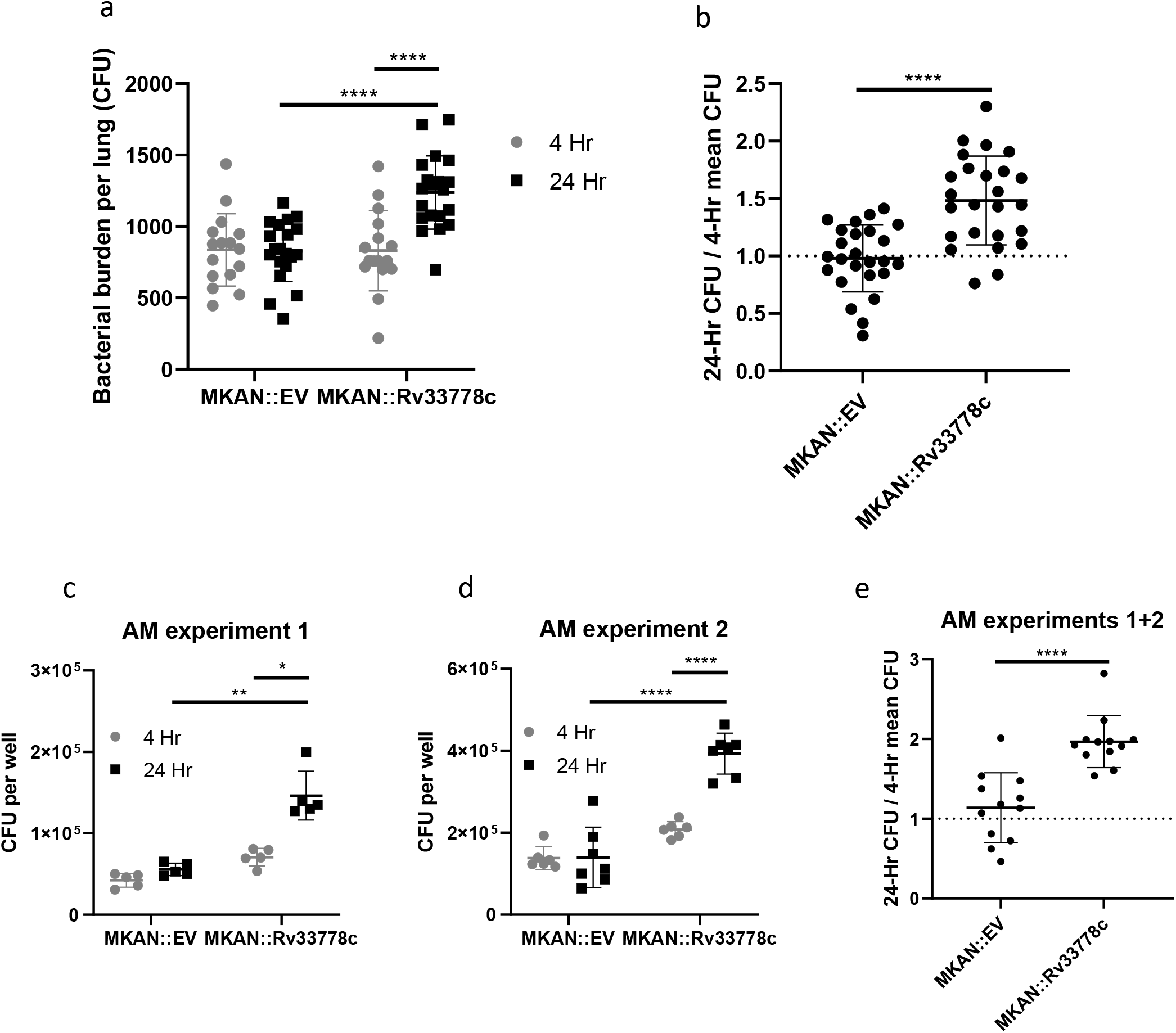
1-TbAd enhances the initial establishment of pulmonary infection. (a-b) CFUs were counted from C57Bl/6 mouse lungs isolated at 4- vs. 24-hours post aerosol infection with *M. kansasii*::EV (MKAN::EV) or *M. kansasii::Rv3377-78c* (MKAN::Rv33778c) (n=20 lungs/condition/time point). (a) Absolute CFU count data are pooled from two independent experiments with similar initial inoculum. (b) 24-hr CFU/mean 4-hr CFU ratio data were pooled from three independent experiments (n=21-25 lungs/condition/time point). (c-e) CFUs were counted from C57Bl/6 murine-derived AMs at 4- and 24-hours after *ex vivo M. kansasii* infection. (c-d) Absolute CFU count data from two independent experiments (N=5 and 7 replicate wells containing infected AMs, respectively, per condition per timepoint). (e) 24-hr CFU/mean 4-hr CFU ratio data pooled from two independent experiments (N=12 per condition). The data are plotted as the mean ± SD. GraphPad Prism 8.1.2 was used to perform Welch’s two-tailed unpaired t-tests where *p<0.05, **p<0.01, ****p<0.0001.

We hypothesized that 1-TbAd was enhancing proliferation of *M. kansasii* in the lungs by promoting survival in resident macrophages. First, using bone marrow-derived macrophages (BMDMs), we infected with *M. kansasii*::EV or *M. kansasii::Rv3377-78c* and collected cell lysates at 4- and 24-hours post infection: the infections proceeded similarly with both bacteria unlike what we had observed in mouse lungs *in vivo* (Sup. Fig. 5). Alveolar macrophages are resident lung macrophages that phagocytose infectious agents entering the lower airways (30). We assessed whether fitness would be altered in an *ex vivo* infection of murine Alveolar macrophages. In two independent experiments, *M. kansasii::Rv3377-78c* increased in numbers by CFU counts from 4 to 24 hours post infection, while *M. kansasii*::EV numbers were largely unchanged and inferior to *M. kansasii::Rv3377-78c* after 24 hours (fig. 6c-d). *M. kansasii::Rv3377-78c* exhibited a 2.0-fold increase compared to the 1.1-fold increase seen in *M. kansasii*::EV (Fig. 6e). This is consistent with our observations in murine lungs and with the conclusion that 1-TbAd provides an advantage to *M. kansasii* during the early stages of pulmonary infection by specifically promoting survival or growth in resident alveolar macrophages.

### 1-TbAd production does not enhance long-term persistence of *M. kansasii* during mixed lung infections

We validated a translaryngeal infection model wherein WT *M. kansasii* was directly introduced into the upper respiratory tract (Sup. Fig. 6 and methods) (31). Mice were monitored for 42 days; bacteria persisted at the same log CFU as the initial infection and the mice did not become overtly sick (Sup. Fig. 6). To test the effect of 1-TbAd in a high-dose competitive infection, wherein we expected high numbers of bacteria to allow us to see subtle changes in bacterial burden, we used the translaryngeal infection model to generate a mixed infection with 3×10^6^ CFUs of a 1:1 WT *M. kansasii* and *M. kansasii::Rv3377-78c* (Fig. 7a). Lung homogenates plated on 7H10 plates with and without 50 μg/ml hygromycin revealed a statistically significant decrease in the *M. kansasii::Rv3377-78c* to WT *M. kansasii* ratio over time, with an initial decrease in the proportion of *M. kansasii::Rv3377-78c* in the bacterial population from 0.48 (week 0) to 0.27 (week 1), stabilizing at that latter proportion over time (Fig. 7a). These findings show no beneficial effect of 1-TbAd production for *M. kansasii* survival *in vivo*.

**Figure 7 –.**
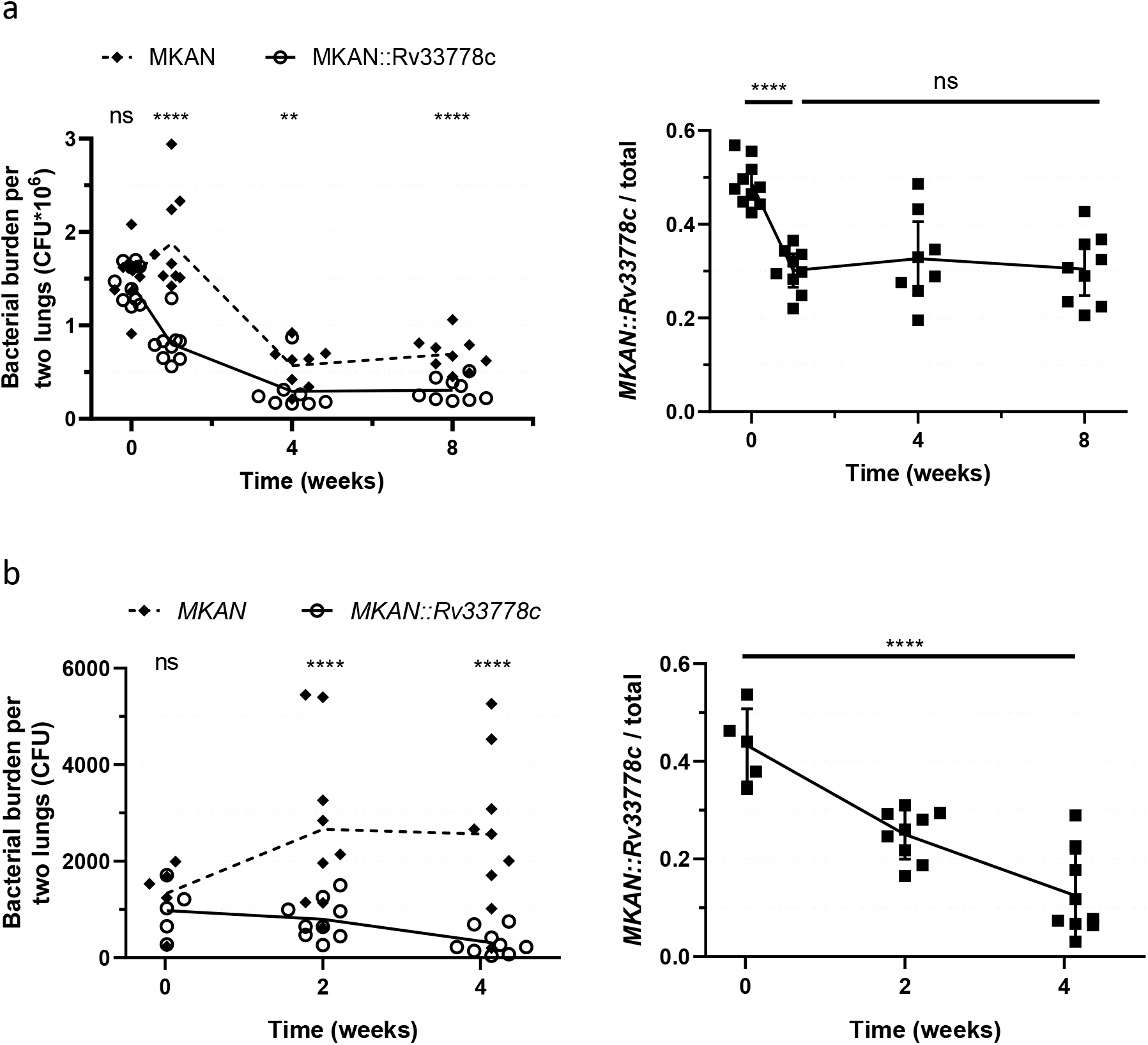
1-TbAd production hinders long-term bacterial growth. (a-b) Mixed 1:1 bacterial suspensions of WT *M. kansasii* (MKAN) and *M. kansasii::Rv3377-78c (MKAN::Rv33778c*) were used to infect (a) WT C57Bl/6 mouse lungs isolated at 1, 7, 28 and 56 days post high-dose trans-laryngeal intubation (n=8-10 lung pairs per timepoint) or (b) *Ccr2 -/-* C57Bl/6 mouse lungs isolated at 0, 14 and 29 days post aerosolization (n=5-9 lung pairs per timepoint). CFUs were counted on 7H10 plates + PANTA ± hyg50. The graphs on the right represent the proportion of *M. kansasii::Rv3377-78c* over the total number of bacteria (MKAN + MKAN::Rv33778c) per mouse per timepoint. The raw data (left) and proportions (right) are plotted as individual datapoints (± SD for proportions only). GraphPad Prism 8.1.2 was used to perform the ratio paired t-test (ratio per timepoint/left) and ordinary one-way ANOVA (proportions over time/right).

C-C chemokine receptor 2 (CCR2) is an essential component for defense in the airways; *Ccr2-/-* mice lose the ability to recruit non-tissue resident immune cells and succumb to *M. tuberculosis* infection (32). We used these mice to test whether the short-term alveolar macrophage phenotype could be recapitulated in a longer-term *in vivo* setting without the interference of recruited immune cells in WT mice that might explain the lack of a phenotype in the previous experiment. We aerosol infected *Ccr2-/-* mice with a mixed 1:1 bacterial suspension of WT *M. kansasii* and *M. kansasii::Rv3377-78c* (Fig. 7b). Interestingly, although WT *M. kansasii* exhibited a slight increase in numbers over time, *M. kansasii::Rv3377-78c* steadily decreased within the same period. A statistically significant decrease in the *M. kansasii::Rv3377-78c* to WT *M. kansasii* ratio over time was noted, with a steady, non-stabilizing decrease in the proportion of *M. kansasii::Rv3377-78c* in the bacterial population (Fig. 7b). With these unexpected results, to be sure that our method of identifying *M. kansasii::Rv3377-78c* (hygromycin resistance) in the mixed infection was valid we compared hygromycin resistance of *M. kansasii*::EV (sup. Fig 7a) and *M. kansasii::Rv3377-78c* (sup. Fig 7b) after four weeks *in vivo* from separate aerosol infections. Hygromycin resistance declined by up to 20% initial levels (sup. Fig 7b); this is clearly less than the 40-70% decline observed in the proportion of *M. kansasii::Rv3377-78c* during mixed infections (Fig 7). Neither strain appeared more fit in the mouse in the separate infection (sup. Fig 7c). Thus, 1-TbAd production clearly did not enhance *M. kansasii* survival in any of these *in vivo* infection models, but on the contrary may hinder fitness in the long term.

## DISCUSSION

Our results indicate the feasibility of using *M. kansasii* to study the pathoevolution of *M. tuberculosis*. The less virulent non-tuberculous mycobacterium (NTM) species is a suitable surrogate for the expression of *M. tuberculosis*-specific products, such as 1-TbAd, and can readily be used *in vitro*, and for *ex vivo* and *in vivo* experimental infection models. We showed that 1-TbAd led to an improved survival during the first 24 hours of infection when tested *in vivo*, and *ex vivo* in alveolar macrophages, but the isolated addition of 1-TbAd to *M. kansasii* resulted in impaired persistence in different murine infection models.

In the current study, we demonstrated that *M. kansasii::Rv3377-78c* produced lipid species distinct from those seen in *M. kansasii*::EV. Our prior and current data indicated that transfer of TbAd biosynthesis genes to *M. kansasii* does not promote growth at neutral pH, but confers increased growth in 7H9 media in the more acidic pH range (5.0-5.4) (22). When added as a pure compound or produced by *M. tuberculosis*, 1-TbAd detectably raises the pH and swells lysosomes in human macrophages (18, 22). In broad terms, the mechanisms by which the effects of 1-TbAd are mediated could occur through direct chemical results of lysosomotropism, or through signalling.

1-TbAd could act as an amphipathic weak base that penetrates membranes as an uncharged conjugate base to selectively accumulate in acidic phagolysosomes where proton capture raises pH and confers a positive charge, trapping the compound and leading to lysosomal swelling. We did not see clear *Rv3377-Rv3378c*-dependent alkalization of 7H9 broth which might be explained by compartmentalization: whereas intracellular bacteria are bound in a small phagosomal compartment of 10^−15^ L, growth in 7H9 media provides a much larger compartment for 1-TbAd to disperse in if it is physically shed from the bacterium. By a rough estimate, 10^10^ bacteria-worth of 1-TbAd would be required to change the pH of 1 ml of 7H9 from 5.2 to 5.3, about 10,000 times the concentration of bacteria we inoculate (see supplementary data calculation). This result suggested that 1-TbAd production provides a pH-dependent growth advantage intrinsic to the bacterium, separate from but not exclusive to lysosomal perturbation.

In phagocyte-free systems, we experimentally tested the hypothesis that the pH-dependent growth advantage of *M. kansasii* producing 1-TbAd might involve direct contact of the molecules with bacteria. Overall we found that direct exposure to externally added, protonated 1-TbAd at low and high concentrations did not promote growth as might be expected of a signaling molecule. Increased growth was selectively observed at low pH (5.1-5.4) when 1-TbAd was generated through gene transfer and the action of enzymes in the bacterium. While not fully understood, these divergent outcomes whereby the compartment of origin controls the protective effect can be explained by the lysosomotropy model. Cytosolic 1-TbAd would be expected to shed its proton during membrane passage into the periplasm. The lack of pH control in the cytosol of *M. kansasii::Rv3377-78c* cultured in acidic media suggests that 1-TbAd does not act as a shield against proton flow into *M. kansasii* cytosol, which is the expected outcome if membrane penetration is required for the protective effect. These outcomes indicate that the consistently observed survival advantage could derive from 1-TbAd passage from the cytosolic membrane into the periplasm, mycolate membrane or surface of *M. kansasii*. 1-TbAd may act on the bacterial population itself by targeting or protecting specific molecules during exposure to low pH, stopping damage from occurring.

It is noteworthy that genetic and chemical complementation provide different information about mycobacterial virulence factors, which in this case might result from the differential compartmentalization of the molecules. This result also argues that 1-TbAd must exert its protective effect at a specific location within the bacterial cell or cell wall. Exogenous 1-TbAd may simply not reach this specific location, or not reach the location in an uncharged conjugate base state. Together with our genetic complementation data, it is clear that *Rv3377c-Rv3378c*-dependent metabolites including 1-TbAd do not have a direct growth-promoting effect. Our data also demonstrate that *Rv3377c-Rv3378c*-dependent metabolites protect against acid stress *in vitro*, using mechanisms that are independent of macrophage function, including lysosomes or activating receptors.

We did not directly characterize the impact of 1-TbAd on the *M. kansasii* cell envelope composition, therefore we can formally assign effects to Rv3377c and Rv3378c but cannot refute the possibility of an indirect pathway for the 1-TbAd effect. It is notable that the overall lipid profiles examined by TLC were not significantly altered by gene transfer. The Congo Red retention assay (25) and the retained ability to produce carotenoid pigments (24) and turn yellow upon light exposure both provide indirect evidence that the overall composition of the cell membrane has been preserved.

Another key finding is that the complemented strain fared better than *M. kansasii*::EV in the initial stages of *in vivo* infection. We subsequently showed that *M. kansasii::Rv3377-78c* was more fit to thrive inside alveolar macrophages, but not BMDMs within that same timeframe, demonstrating that the production of 1-TbAd can subvert the first lines of host defense encountered by the pathogen This finding is in line with recent findings describing differential replication potentials for *M. tuberculosis* in BMDMs vs. alveolar macrophages, with the latter being more permissive than IFN-⍰- or LPS-activated BMDMs for *M. tuberculosis* replication (33). It is important to note that the BMDMs used to assess *M. kansasii::Rv3377-78c* were not activated with IFN-⍰ or LPS. Future work focused alternatively on the host will be needed to characterize what fundamental differences between different cell types, in different activation states, play a role in the 1-TbAd response.

WT *M. kansasii* appeared to have outcompeted *M. kansasii::Rv3377-78c* in low- and high-dose mixed infection settings in both WT and *Ccr2-/-* mice. Therefore, although expression of *Rv3377c-Rv33778c* conferred a survival benefit to *M. kansasii* in specific *in vitro* and short-term infection contexts, there may be a drawback to 1-TbAd expression in the non-adapted mycobacterium for persisting in the murine host. The decrease in proportion of *M. kansasii::Rv3377-78c* with mixed infection is not entirely explained by functional loss of hygromycin resistance over time *in vivo*, strictly according to our numerical data. We hypothesize that the burden of constitutive production of 1-TbAd, which sequesters adenosine molecules, may prevent energy storage in the form of ATP and have a negative impact on long-term *in vivo* survival for *M. kansasii*. Another consideration is the extent to which mycobacterial killing is dependent on acid-mediated mechanisms. The intrinsic antacid properties of 1-TbAd, its tropism for acid compartments, its marked remodeling of lysosomes and the pH-dependent basis of growth promotion in culture media all point at a selective role in protection against acid-mediated killing. Therefore, the varied outcomes in the models examined herein might depend on the extent to which they test acid-dependent killing. One question that remains unanswered is whether there is a single, predominant mechanism of action for which 1-TbAd production is mainly conserved in *M. tuberculosis*, or multiple important functions.

Phenotypes observed after pathogen-specific genetic complementation into non-pathogenic species provide different information than the more commonly observed loss-of-function phenotypes observed after deleting genes from pathogens. The latter requires breaking one link in a causal chain and might have rippling downstream effects, if it is not the final component of a response cascade. Gain of function is a rare phenomenon that occurs only when the components of a larger pathogen-specific system are fully recapitulated in the non-pathogen and then tested under conditions in which this system is essential. In this regard, that biosynthetic genes for 1-TbAd can promote early stage growth in alveolar macrophages was unexpected, so these data now point to a new direction for mechanistic studies of these genes in *M. tuberculosis*.

To date, the established virulence factors of *M. tuberculosis* are largely conserved among the NTM, with the exception of a few proteins and lipids like 1-TbAd, Tuberculosis Necrotizing Toxin (34) and the MoaA1-4 operon (35), aprABC (36) and others. Our data indicate that 1-TbAd alone does not confer a long-term *in vivo* benefit consistent with *in vitro* and *in vivo* phenotypic differences between *M. tuberculosis* and *M. kansasii*. Therefore, events of acquisition of other *M. tuberculosis*-specific and loss of *M. kansasii*-specific effectors are likely required to recreate an *M. tuberculosis*-like *M. kansasii* mutant strain, or alternatively the difference is due to the compounding of multiple subtle effects that complement one another. The possibility that mycobacterial virulence factors manifest their phenotype in a cell-dependent fashion is consistent with the known transcellular lifestyle of *M. tuberculosis* and suggests that different host-cell types should be used to detect undiscovered virulence determinants.

Viewed in this light, using genetically complemented NTM is therefore useful to single out the effects of specific elements that contributed to *M. tuberculosis* host adaptation without producing a clearly hypervirulent NTM. The 1-TbAd family of molecules represents a newly discovered pathogen-specific collection of compounds that has no clear chemical analog in other bacterial systems, and their exact mechanisms of action remain elusive. We can conclude from our study that *Rv3377c-Rv3378c* transfer acts in a eukaryotic cell-free system by a localized chemical mechanism that involves pH, and that such changes can be determinative of outcomes in alveolar macrophages at expectedly early time points post-infection. As such, these molecules may aid in the establishment of infection within the lower respiratory tract.

## MATERIALS AND METHODS

### Bacterial strains and culture conditions

*M. kansasii* ATCC 12478 and *M. tuberculosis* H37Rv were grown in Middlebrook 7H9 broth (BD Difco, MD, USA) as previously described (3). To test the ability of *M. kansasii::Rv3377-78c* to produce yellow pigment, fully-formed colonies were additionally exposed to white light and incubated at room temperature for 7 days. Where indicated and to ensure single-cell suspensions, liquid bacterial cultures were de-clumped by slowly passaging through 5x 22G, 5x 25-G and 3x 26-G needles followed by low-speed centrifugation at 50 g for 5 minutes with passage through a 5-μm filter. To generate *M. kansasii::Rv3377-78c*, a 2.4-kb PCR fragment spanning *Rv3377c-Rv3378c* was generated using primers BamHI-*Rv3377c-Rv3378c-F* and HindIII-*Rv3377c-Rv3378c*-R (Sup. Table 1) using high-fidelity Phusion DNA polymerase (New England Biolabs). The fragment was subsequently digested with BamHI and HindIII (all restriction enzymes from New England Biolabs) and cloned into the episomal plasmid pMV261 with the constitutive mycobacterial *hsp60* promoter and a selective apramycin resistance marker. The *hsp60-Rv3377c-Rv3378c* fragment was shuttled into the integrative vector pMV306 containing a hygromycin resistance cassette using XbaI and HindIII. All ligations were done using T4 DNA ligase (Fermentas). The resulting plasmid pMV306::*Hsp60-Rv3377c-Rv3378c* was verified by Sanger sequencing (Genome Québec) to ensure the absence of frameshift or point mutations during the cloning process. An unaltered version of pMV306 with a hygromycin resistance cassette was used to create the empty vector control strain *M. kansasii*::EV. Following electroporation, *M. kansasii*::EV and *M. kansasii::Rv3377-78c* were grown in the presence of 100 μg/mL hygromycin (Wisent).

### Detection of cell filtrate adenosine-linked lipids

*M. kansasii*::EV, *M. kansasii::Rv3377-78c* and *M. tuberculosis* were grown to mid-log phase and subsequently incubated with 0.25 μCi/mL radiolabeled [8-^14^C]adenosine (American Radiolabeled Chemicals) for 14 days. Polar lipid fractions were extracted using CHCl_3_:CH_3_OH:0.3%NaCl (v/v/v)(22).)(37). Extracted lipids were spotted on a TLC Silica Gel 60 (Millipore Sigma) with CHCl_3_:CH_3_OH:H_2_O 10:5:1 (v/v/v) used as the mobile phase solvent. The radiolabeled signature was developed using Storm 840 PhosphorImager (GE Healthcare) to visualize adenosine-linked lipids in each lane. [8-^14^C]adenosine 1:100 was used as a no-lipid staining control. The plate was stained with 5% phosphomolybdic acid reagent (PMA) (Sigma) and heated briefly using an industrial blow-dryer to visualize the total amounts of lipids loaded in each lane.

### HPLC-MS analysis of lipids from cells and supernatant

*M. kansasii::Rv3377-78c, M. kansasii*::Empty vector (EV) and *M. kansasii* parent strain were grown in 30 ml of 7H9 media supplemented with albumin-dextrose-saline (5% Bovine Serum Albumin Fraction V, 2% anhydrous dextrose and 0.87% sodium chloride) to late log-phase. Bacterial cell pellet and supernatant were separated by centrifugation at 5000 rpm for 5 minutes. The cell pellet was resuspended in 4 ml of PBS at pH 7.4 and distributed equally into four 2 ml screw-cap tubes. The cells were pelleted by centrifugation and further resuspended in 1 ml PBS at pH titrated to 7.4, 6.4, 5.5 and 4.5 with hydrochloric acid and incubated for 2 hours at 37°C. At the end of 2 hours the cell pellet and the PBS supernatant were collected for lipid extraction. Added 10 volumes (3 ml) of Chloroform/methanol (C/M) at the ratio of 1:2 and extracted for 1 hour at room temperature. A second extraction under similar conditions was performed with 3 ml of C/M at the ratio of 1:1. The extracted fractions were pooled and dried under a stream of nitrogen gas. Lipids from the 1 ml PBS supernatant was extracted using acidified ethyl acetate by adding 3 μl of 6N HCl and 1.4 ml of ethyl acetate and mixing for 30 minutes in an Orbitron shaker. The mixture was centrifuged at 2000 rpm for 15 minutes to collect the upper organic phase and dried on to glass under a stream of nitrogen gas at room temperature, and total lipids were weighed using analytical balance. HPLC-MS separations were performed as described (17) using equal amount of lipid samples from different experimental conditions as determined by weight on a Mettler balance.

### Congo Red uptake assay

Bacterial cultures were grown on Congo Red-containing 7H10 plates for 14 days, scraped into a 15-ml conical tube, washed with water until the supernatant became clear and incubated with 2 ml DMSO for 2 hours (25, 38). Congo Red was measured in the resultant supernatant at A488. The values were normalized to the dry weight of the pellet to define the Congo Red binding index.

### Extracellular pH measurement

For all pH experiments, liquid media was prepared as usual and the pH was equilibrated to 4.0, 4.9, 5.0, 5.1, 5.2, 5.4, 6.0, 6.7 and 7.2 using 2 M HCl or NaOH. OD_600_ was adjusted to 0.34 and 222 μl of mid-log phase de-clumped bacteria were added to 15 ml of freshly prepared, pH-adjusted 7H9 in 150-ml roller bottles (final OD_600_ of 0.005). Triplicate cultures were made per condition (strain and pH) and incubated at 37°C, rolling in the dark. OD_600_ was measured every 2-3 days using 2 x 200 μl of culture (technical duplicates) and a Tecan Infinite M200 Pro plate reader. At days 8 and 17, 1 ml was removed from each culture for centrifugation and recovery of supernatant, which was stored at 4°C until extracellular pH was read using micro pH combination electrode (AgCl) (Sigma Aldrich), and Orion Star A111 meter (Thermo Scientific).

### Synthetic TbAds and chemical complementation

Synthetic 1-TbAd and *N*^6^-TbAd were produced as described previously (39). Bacterial cultures were grown to mid-log phase and de-clumped as described above, then inoculated into 96-well plates in 200 μl pH-adjusted 7H9 containing 1-TbAd, *N*^6^-TbAd or vehicle (DMSO) control as indicated. Plates were incubated at 37°C in the dark and OD_600_ was measured every one to three days with a Tecan Infinite M200 Pro plate reader.

### Intracellular pH measurement

Bacterial cultures were grown to mid-log phase and de-clumped as previously described. 5×10^8^ CFU were pelleted, the supernatant completely removed, and the cells resuspended in 0.3 ml PBS containing 100 μM Carboxyfluorescein diacetate succinimidyl ester (CFDA-SE) (CellTrace™ CFSE, ThermoFisher) for 20 minutes at 37°C, shaking at 150 RPM in the dark. Bacteria were next diluted with 10 ml of 7H9 and incubated for 4 hours at 37°C, rolling in the dark. A portion of bacteria was taken during this incubation for lysis in normal saline (0.9% NaCl) by beating with silica beads (MP Biomedicals, FastPrep-24) to extract free protein-CFSE conjugate to generate a fluorescence-to-pH standard curve. After 4 hours, bacteria were washed and resuspended in normal saline to an OD_600_ of 0.4. 20 μl of bacteria in saline were sub-cultured into 180 μl of freshly prepared, pH-adjusted 7H9 in 96-well plates (opaque-black for fluorescence (ThermoScientific Nunclon Delta Surface), translucent-colourless for absorbance (Falcon). Lysed bacteria were plated similarly for the pH standard curve. Immediately, plates were placed in plate readers (Tecan Infinite M200 Pro) at 37°C, shaking and measuring fluorescence or OD_600_ every 30 minutes. 528-nm fluorescence was measured from 490-nm excitation (pH-sensitive), and 520-nm fluorescence was measured from 450-nm excitation (pH-insensitive). To calculate pH, 7H9 background was subtracted for all data first (pH did not alter 7H9 fluorescence). Next, a standard curve of 490-excitation/450-excitation in relation to 7H9 pH was created from the CFSE-containing cell lysate. The 490/450 ratios calculated from the culture wells were applied to the standard curve to determine intracellular pH.

### Murine pulmonary infection

Male and female C57Bl/6 and *Ccr2-/-* mice (Jackson Laboratories) were used for experiments. Mice were approximately 6-16 weeks of age upon infection over all experiments; different groups were age- and sex-matched. All protocols were approved by independent ethics oversight at the RI-MUHC and followed the guidelines of the Canadian Council on Animal Care (CCAC). C57Bl/6 mice were infected through aerosolization (ONARES, NJ, USA) of bacterial cultures at OD_600_ 0.4 for 15 minutes as previously described (3). Alternatively, C57Bl/6 mice were infected via trans-laryngeal intubation using 50 μl of a high-dose mixed bacterial suspension containing both wild-type (WT) *M. kansasii* and *M. kansasii::Rv3377-78c* at OD_600_ 0.2. C57Bl/6 *Ccr2-/-* (C-C motif chemokine receptor 2 knockout) mice were infected through aerosolization of a low-dose mixed suspension at OD_600_ 1.0. Mouse lungs were harvested at 4 or 24 hours (early time-points to measure short-term bacterial establishment) and 14, 21, 28, 42 or 56 days post infection (later time-points to measure long-term bacterial persistence) into 1ml 7H9 and homogenized using an Omni Tissue Homogenizer TH (Omni International) at high speed for 45 seconds. Serial dilutions made in 7H9 liquid media from lung homogenates were plated on 7H10 plates containing PANTA ± 50 μg/ml hygromycin. Colony-forming units (CFU) were counted 2 weeks post-plating to determine bacterial burden.

### Murine macrophage isolation

Bone marrow was isolated from C57Bl/6 murine tibiae and femora. Bone marrow-derived macrophage (BMDMs) were differentiated with recombinant M-CSF (100 U/ml) (Peprotech) for a period of 7 days as previously described (40), after which they were lifted using 4 ml CellStripper Solution (Corning) and seeded into the appropriate tissue-culture plates. For alveolar macrophage (AM) isolation, the respiratory tract including the trachea and lungs was isolated and repeatedly perfused with cold sterile phosphate-buffered saline (PBS) to collect the cells through bronchoalveolar lavage (BAL). Alveolar macrophages were enriched through adherence purification to tissue culture-treated 96-well plates over 24 hours, at which point other cells were washed away. All mammalian cells were cultured in RPMI 1640 media supplemented with non-essential amino acids, 10 mM HEPES, 10% FBS ± Penicillin/Streptomycin (Wisent).

### Macrophage infection

Macrophages were seeded into 96-well plates (100,000 cells/200 μl complete RPMI media without antibiotics). Bacterial cultures were grown to OD_600_ 0.2-0.5, clumps were removed to ensure single-cell suspensions and adjusted in complete RPMI media (without antibiotics) to an OD_600_ 0.01. Macrophages were infected by replacing the media with fresh media containing bacterial suspension. After 4 hours of infection, the wells were gently washed three times with PBS to remove extracellular bacteria and fresh complete RPMI media (without antibiotics) was added to each well. At indicated time points, the plates were spun down at 2,000 g for 5 minutes. Each well was subjected to PBS containing 1% Triton X-100 for 10 minutes at room temperature to induce macrophage lysis. Following serial dilution and plating, CFUs were counted on 7H10 plates 2 weeks post-plating to determine bacterial burden.

### Statistical analysis

All calculations and statistical analyses were performed using Microsoft Excel or GraphPad Prism. Calculations included (1) the ratio of individual 24-hr CFU values/mean 4-hr CFU for murine lung and macrophage infection assays to determine bacterial proliferation [datapoint = (24-hr CFU)/(mean 4-hr CFU)] and (2) the proportion of *MKAN::Rv33778c* / total (values paired from individual mice) in competition assays to determine comparative fitness of WT *M. kansasii* vs. *M. kansasii::Rv3377-78c* [datapoint = (CFU on 7H10-Hygromycin)/(CFU on 7H10)].

## Supporting information

Supplemental Calculation

## Acknowledgements

We would like to thank Dr. Maziar Divangahi and Laura Mendon⍰a for transferring their knowledge on alveolar macrophages and for providing *Ccr2-/-* mice. We would also like to thank Dr. David Young for assistance with chromatography and figure preparation.

This project was funded through the support of an operating CIHR grant FND-148362.

## SUPPLEMENTAL MATERIAL

**Supplementary Table 1** – Primer list

**Supplementary Figure 1 – 1-TbAd production enhances growth at low pH.** *M. kansasii*::EV (MKAN::EV) and *M. kansasii::Rv3377-78c* (MKAN::Rv33778c) cultures were inoculated at equal OD_600_ into fresh pH-adjusted 7H9 (using HCl titration) and incubated at 37°C in a rolling incubator over 17 days. The starting pH of the cultures is indicated above each graph. (a) OD_600_ was monitored every 2-3 days. The data are plotted separately for each of three independently growing cultures per strain. The data are representative of 5 independent experiments.

**Supplementary Figure 2 – Chemical complementation with TbAd does not change 7H9 broth pH nor enhance growth**. (a) pH of 7H9 of *M. kansasii* cultures taken one hour after addition of 20 μM TbAd (or 0.2% DMSO control) measured by addition of fluorescein and reading fluorescence against a standard curve. (b) Additional data for fig. 4 on cultures grown in 7H9 broth set to lower pH (absorbance values are near background).

**Supplementary Figure 3 – *M. kansasii*::EV and *M. kansasii::Rv3377-78c* infections progress similarly in C57Bl/6 mice.** CFUs were counted from C57Bl/6 mouse lungs isolated at 1-, 21- and 42-days post infection (n=5 lungs/condition/time point). The data are plotted as the median.

**Supplementary Figure 4 – *M. kansasii* fitness in murine lungs in the first 24 hours after infection.** CFUs were counted from C57Bl/6 mouse lungs isolated at 4- vs. 24-hours post aerosol infection with *M. kansasii* (n=5-10 lungs/condition/time point). Top row, absolute CFU count data from independent experiments. Bottom row, 24-hr CFU/mean 4-hr CFU ratio data from corresponding independent experiments. The data are plotted as the mean ± SD. GraphPad Prism 8.1.2 was used to perform Welch’s two-tailed unpaired t-tests where ns = not significant (p>0.05), *p<0.05, **p<0.01, ****p<0.0001.

**Supplementary Figure 5 – *M. kansasii*::EV and *M. kansasii::Rv3377-78c* infections progress similarly in BMDMs.** CFUs were counted from C57Bl/6 murine-derived BMDMs infected with *M. kansasii* 4- and 24-hours post-infection. Top, absolute CFU count data are plotted as the mean of technical replicates (N=3, 5 and 7 respectively) ± SD, individually for three independent experiments. Bottom, 24-hr CFU/mean 4-hr CFU ratio data is plotted for individual experiments and pooled (N=15 per condition), shown as the mean ± SD. GraphPad Prism 8.1.2 was used to perform Welch’s two-tailed unpaired t-tests (ns, not significant p>0.05; *p<0.05).

**Supplementary Figure 6 - high-dose infection with *M. kansasii* persists but does not cause debilitating disease in C57Bl/6 mice.** *M. kansasii* was used to infect C57Bl/6 mice with 10^6^ and 10^5^ CFUs. (a) Mice were sacrificed at 1- and 42-days post infection to establish initial and persistent infectious dose, respectively. (b) Mice were weighed over the 42-day period to assess change in weight as a proxy for clinical status.

**Supplementary Figure 7 - *M. kansasii*::EV and *M. kansasii::Rv3377-78c* retain hygromycin resistance in C57Bl/6 mice.** Suspensions of *M. kansasii*::EV (MKAN::EV, panel a) and *M. kansasii::Rv3377-78c (MKAN::Rv33778c*, panel b) were used to infect WT C57Bl/6 mice. Lungs were isolated at 4-hours and 4-weeks post aerosolization (n=5-10 lung pairs per timepoint). CFUs were counted on 7H10 plates + PANTA ± hyg50. a-b, mean pulmonary CFUs determined from plating with or without hyg (solid bars), and percent hyg resistance (+hyg/-hyg x 100%) (empty bars); points represent data from one mouse and bars denote group mean. c, ratio of total pulmonary CFUs of 4 weeks over 4 hours; GraphPad Prism 8.1.2 was used to perform Welch’s two-tailed unpaired t-tests where ns = not significant (p>0.05).

**Supplementary Calculation** – Estimation of 1-TbAd amount required to alter 7H9 pH

## References

1. Peirs P, Lefevre P, Boarbi S, Wang XM, Denis O, Braibant M, Pethe K, Locht C, Huygen K, Content J. 2005. Mycobacterium tuberculosis with disruption in genes encoding the phosphate binding proteins PstS1 and PstS2 is deficient in phosphate uptake and demonstrates reduced in vivo virulence. Infect Immun 73:1898–902.

2. Dunphy KY, Senaratne RH, Masuzawa M, Kendall LV, Riley LW. 2010. Attenuation of Mycobacterium tuberculosis functionally disrupted in a fatty acyl-coenzyme A synthetase gene fadD5. J Infect Dis 201:1232–9.

3. Wang J, McIntosh F, Radomski N, Dewar K, Simeone R, Enninga J, Brosch R, Rocha EP, Veyrier FJ, Behr MA. 2015. Insights on the emergence of Mycobacterium tuberculosis from the analysis of Mycobacterium kansasii. Genome Biol Evol 7:856–70.

4. Bloch KC, Zwerling L, Pletcher MJ, Hahn JA, Gerberding JL, Ostroff SM, Vugia DJ, Reingold AL. 1998. Incidence and clinical implications of isolation of Mycobacterium kansasii: results of a 5-year, population-based study. Ann Intern Med 129:698–704.

5. Johnson MM, Odell JA. 2014. Nontuberculous mycobacterial pulmonary infections. J Thorac Dis 6:210–20.

6. Sheu LC, Tran TM, Jarlsberg LG, Marras TK, Daley CL, Nahid P. 2015. Non-tuberculous mycobacterial infections at San Francisco General Hospital. Clin Respir J 9:436–42.

7. Ricketts WM, O’Shaughnessy TC, van Ingen J. 2014. Human-to-human transmission of Mycobacterium kansasii or victims of a shared source? Eur Respir J 44:1085–7.

8. Griffith DE, Aksamit T, Brown-Elliott BA, Catanzaro A, Daley C, Gordin F, Holland SM, Horsburgh R, Huitt G, Iademarco MF, Iseman M, Olivier K, Ruoss S, von Reyn CF, Wallace RJ, Jr., Winthrop K, Subcommittee ATSMD, American Thoracic S, Infectious Disease Society of A. 2007. An official ATS/IDSA statement: diagnosis, treatment, and prevention of nontuberculous mycobacterial diseases. Am J Respir Crit Care Med 175:367–416.

9. Fortune SM, Jaeger A, Sarracino DA, Chase MR, Sassetti CM, Sherman DR, Bloom BR, Rubin EJ. 2005. Mutually dependent secretion of proteins required for mycobacterial virulence. Proc Natl Acad Sci U S A 102:10676–81.

10. Boon C, Dick T. 2002. Mycobacterium bovis BCG response regulator essential for hypoxic dormancy. J Bacteriol 184:6760–7.

11. Walters SB, Dubnau E, Kolesnikova I, Laval F, Daffe M, Smith I. 2006. The Mycobacterium tuberculosis PhoPR two-component system regulates genes essential for virulence and complex lipid biosynthesis. Mol Microbiol 60:312–30.

12. Salah IB, Ghigo E, Drancourt M. 2009. Free-living amoebae, a training field for macrophage resistance of mycobacteria. Clin Microbiol Infect 15:894–905.

13. Veyrier F, Pletzer D, Turenne C, Behr MA. 2009. Phylogenetic detection of horizontal gene transfer during the step-wise genesis of Mycobacterium tuberculosis. BMC Evol Biol 9:196.

14. Becq J, Gutierrez MC, Rosas-Magallanes V, Rauzier J, Gicquel B, Neyrolles O, Deschavanne P. 2007. Contribution of horizontally acquired genomic islands to the evolution of the tubercle bacilli. Mol Biol Evol 24:1861–71.

15. Boritsch EC, Khanna V, Pawlik A, HonorÈ N, Navas VH, Ma L, Bouchier C, Seemann T, Supply P, Stinear TP, Brosch R. 2016. Key experimental evidence of chromosomal DNA transfer among selected tuberculosis-causing mycobacteria. Proceedings of the National Academy of Sciences of the United States of America 113:9876–81.

16. Sapriel G, Brosch R. 2019. Shared pathogenomic patterns characterize a new phylotype, revealing transition towards host-adaptation long before speciation of Mycobacterium tuberculosis. Genome Biol Evol doi:10.1093/gbe/evz162.

17. Layre E, Lee HJ, Young DC, Martinot AJ, Buter J, Minnaard AJ, Annand JW, Fortune SM, Snider BB, Matsunaga I, Rubin EJ, Alber T, Moody DB. 2014. Molecular profiling of Mycobacterium tuberculosis identifies tuberculosinyl nucleoside products of the virulence-associated enzyme Rv3378c. Proc Natl Acad Sci U S A 111:2978–83.

18. Young DC, Layre E, Pan SJ, Tapley A, Adamson J, Seshadri C, Wu Z, Buter J, Minnaard AJ, Coscolla M, Gagneux S, Copin R, Ernst JD, Bishai WR, Snider BB, Moody DB. 2015. In vivo biosynthesis of terpene nucleosides provides unique chemical markers of Mycobacterium tuberculosis infection. Chem Biol 22:516–526.

19. Mann FM, Xu M, Chen X, Fulton DB, Russell DG, Peters RJ. 2009. Edaxadiene: a new bioactive diterpene from Mycobacterium tuberculosis. J Am Chem Soc 131:17526–7.

20. Lau SK, Lam CW, Curreem SO, Lee KC, Lau CC, Chow WN, Ngan AH, To KK, Chan JF, Hung IF, Yam WC, Yuen KY, Woo PC. 2015. Identification of specific metabolites in culture supernatant of Mycobacterium tuberculosis using metabolomics: exploration of potential biomarkers. Emerg Microbes Infect 4:e6.

21. David HL. 1974. Carotenoid pigments of Mycobacterium kansasii. Appl Microbiol 28:696–9.

22. Buter J, Cheng TY, Ghanem M, Grootemaat AE, Raman S, Feng X, Plantijn AR, Ennis T, Wang J, Cotton RN, Layre E, Ramnarine AK, Mayfield JA, Young DC, Jezek Martinot A, Siddiqi N, Wakabayashi S, Botella H, Calderon R, Murray M, Ehrt S, Snider BB, Reed MB, Oldfield E, Tan S, Rubin EJ, Behr MA, van der Wel NN, Minnaard AJ, Moody DB. 2019. Mycobacterium tuberculosis releases an antacid that remodels phagosomes. Nat Chem Biol 15:889–899.

23. de Duve C, Trouet A, Campeneere DD, Baurian R. 1978. Liposomes as lysosomotropic carriers. Ann N Y Acad Sci 308:226–34.

24. Kirti K, Amita S, Priti S, Mukesh Kumar A, Jyoti S. 2014. Colorful World of Microbes: Carotenoids and Their Applications. Advances in Biology 2014:837891.

25. Jankute M, Nataraj V, Lee OY, Wu HHT, Ridell M, Garton NJ, Barer MR, Minnikin DE, Bhatt A, Besra GS. 2017. The role of hydrophobicity in tuberculosis evolution and pathogenicity. Sci Rep 7:1315.

26. Portaels F, Pattyn SR. 1982. Growth of mycobacteria in relation to the pH of the medium. Ann Microbiol (Paris) 133:213–21.

27. Vandal OH, Nathan CF, Ehrt S. 2009. Acid resistance in Mycobacterium tuberculosis. J Bacteriol 191:4714–21.

28. Pethe K, Swenson DL, Alonso S, Anderson J, Wang C, Russell DG. 2004. Isolation of Mycobacterium tuberculosis mutants defective in the arrest of phagosome maturation. Proc Natl Acad Sci U S A 101:13642–7.

29. Stewart GR, Patel J, Robertson BD, Rae A, Young DB. 2005. Mycobacterial mutants with defective control of phagosomal acidification. PLoS Pathog 1:269–78.

30. Johnson JD, Hand WL, King NL, Hughes CG. 1975. Activation of alveolar macrophages after lower respiratory tract infection. J Immunol 115:80–4.

31. Guilbault C, Martin P, Houle D, Boghdady ML, Guiot MC, Marion D, Radzioch D. 2005. Cystic fibrosis lung disease following infection with Pseudomonas aeruginosa in Cftr knockout mice using novel non-invasive direct pulmonary infection technique. Laboratory Animals 39:336–352.

32. Peters W, Scott HM, Chambers HF, Flynn JL, Charo IF, Ernst JD. 2001. Chemokine receptor 2 serves an early and essential role in resistance to Mycobacterium tuberculosis. Proc Natl Acad Sci U S A 98:7958–63.

33. Huang L, Nazarova EV, Tan S, Liu Y, Russell DG. 2018. Growth of Mycobacterium tuberculosis in vivo segregates with host macrophage metabolism and ontogeny. The Journal of experimental medicine 215:1135–1152.

34. Pajuelo D, Gonzalez-Juarbe N, Tak U, Sun J, Orihuela CJ, Niederweis M. 2018. NAD(+) Depletion Triggers Macrophage Necroptosis, a Cell Death Pathway Exploited by Mycobacterium tuberculosis. Cell Rep 24:429–440.

35. Levillain F, Poquet Y, Mallet L, Mazeres S, Marceau M, Brosch R, Bange FC, Supply P, Magalon A, Neyrolles O. 2017. Horizontal acquisition of a hypoxia-responsive molybdenum cofactor biosynthesis pathway contributed to Mycobacterium tuberculosis pathoadaptation. PLoS Pathog 13:e1006752.

36. Abramovitch RB, Rohde KH, Hsu F-F, Russell DG. 2011. aprABC: a Mycobacterium tuberculosis complex-specific locus that modulates pH-driven adaptation to the macrophage phagosome. Molecular microbiology 80:678–694.

37. Slayden RA, Barry CE, 3rd. 2001. Analysis of the Lipids of Mycobacterium tuberculosis. Methods Mol Med 54:229–45.

38. Cangelosi GA, Palermo CO, Laurent JP, Hamlin AM, Brabant WH. 1999. Colony morphotypes on Congo red agar segregate along species and drug susceptibility lines in the Mycobacterium avium-intracellulare complex. Microbiology 145 (Pt 6):1317–24.

39. Buter J, Heijnen D, Wan IC, Bickelhaupt FM, Young DC, Otten E, Moody DB, Minnaard AJ. 2016. Stereoselective Synthesis of 1-Tuberculosinyl Adenosine; a Virulence Factor of Mycobacterium tuberculosis. J Org Chem 81:6686–96.

40. Zhang X, Goncalves R, Mosser DM. 2008. The isolation and characterization of murine macrophages. Curr Protoc Immunol Chapter 14:Unit 14 1.

