## Supplemental Calculation for "Heterologous production of 1-tuberculosinyladenosine in *Mycobacterium kansasii* models pathoevolution towards the transcellular lifestyle of *Mycobacterium tuberculosis*"

Estimation for amount of *M. kansasii::Rv3377-78c* required to produce enough 1-TbAd to detectably raise the pH of 1 ml of 7H9 :

Assumptions:

- negligible buffering from 7H9
- pH change from 5.2 to 5.3
- Every 1-TbAd molecule can capture one proton (no intermediate equilibrium)
- *M. tuberculosis* contains up to 7 x 10^-17 g of 1-TbAd in one cell (22); assume *M. kansasii::Rv3377-78c* contains the same amount and that all of it is available for neutralization.

1-TbAd is 540 g/mol. Therefore, one cell contains 1.30 x 10^-19 mol 1-TbAd. (alternatively put, 1.30 x 10^-19 mol x 6.02 x 10^23 molecules/mol = 78,300 molecules of 1-TbAd per cell). For a pH change of 5.2 to 5.3, the difference in number of H+ ions is (10^-5.3 M) - (10^-5.2 M) = 1.30 x 10^-6 M of H+. For 1 ml of solution, that is 1.30 x 10^-9 moles of H+. Therefore, the number of bacteria needed to change the pH from 5.2 to 5.3 in 1 ml is (1.30 x 10^-9mol) / (1.30 x 10^-19 mol/cell) = 10^10 bacteria. This is an $OD_{600}$ of roughly 100.
