## Supplementary figures and images for "Heterologous production of 1-tuberculosinyladenosine in *Mycobacterium kansasii* models pathoevolution towards the transcellular lifestyle of *Mycobacterium tuberculosis*"

### Supplemental Figure 1

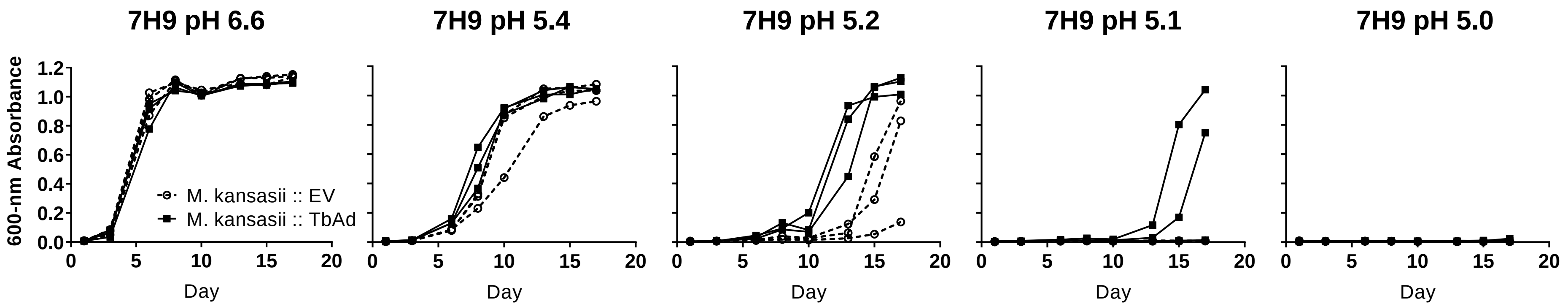

### Supplemental Figure 2

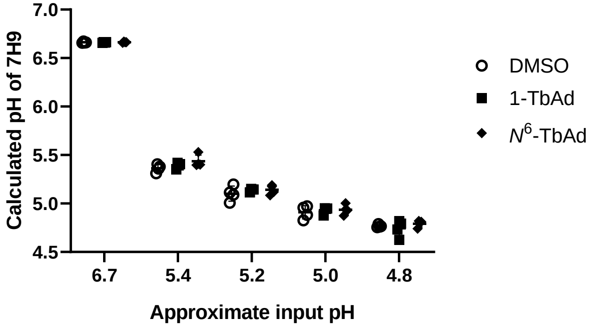

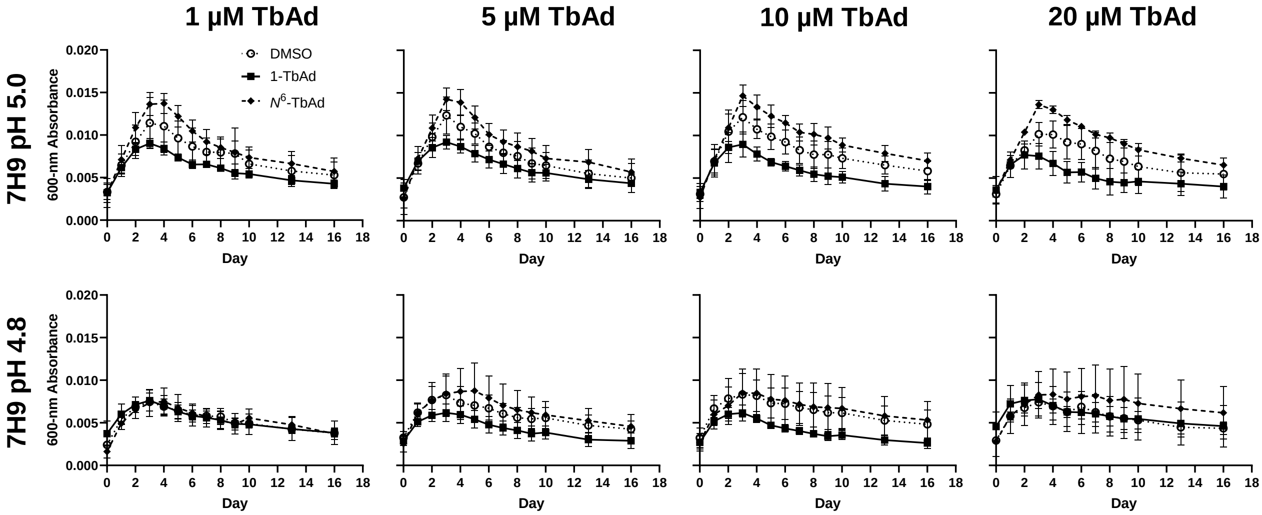

### Supplemental Figure 3

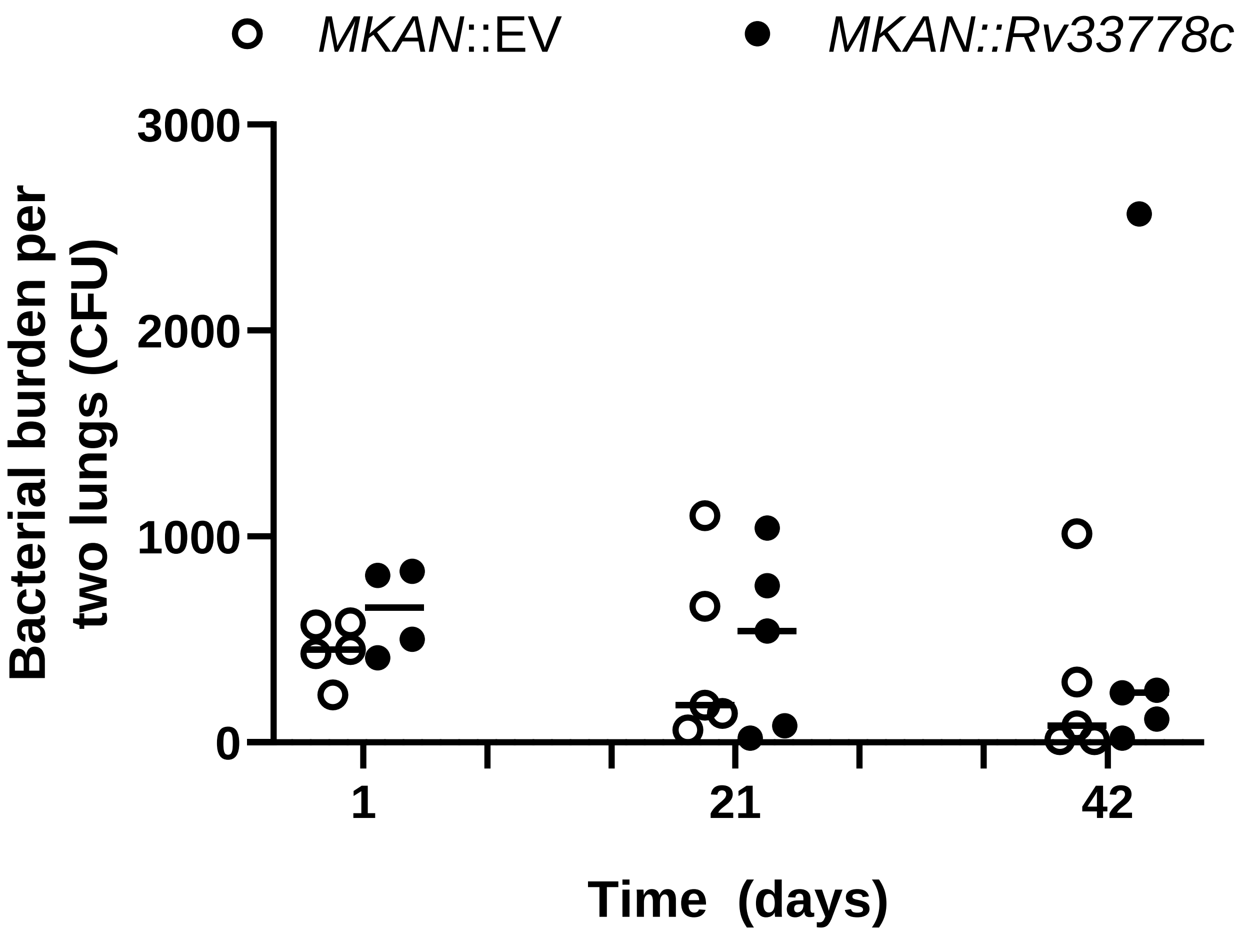

### Supplemental Figure 4

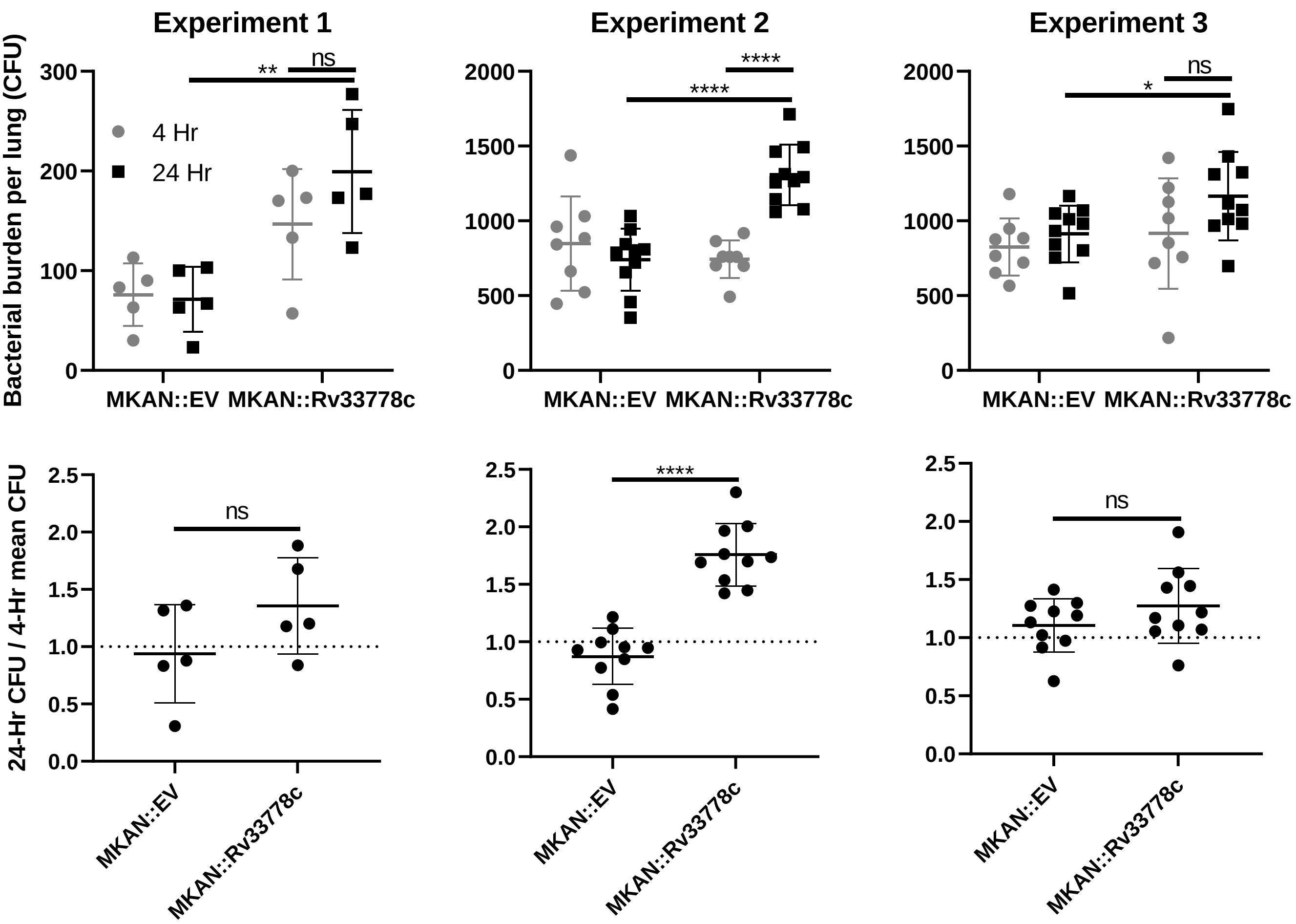

### Supplemental Figure 5

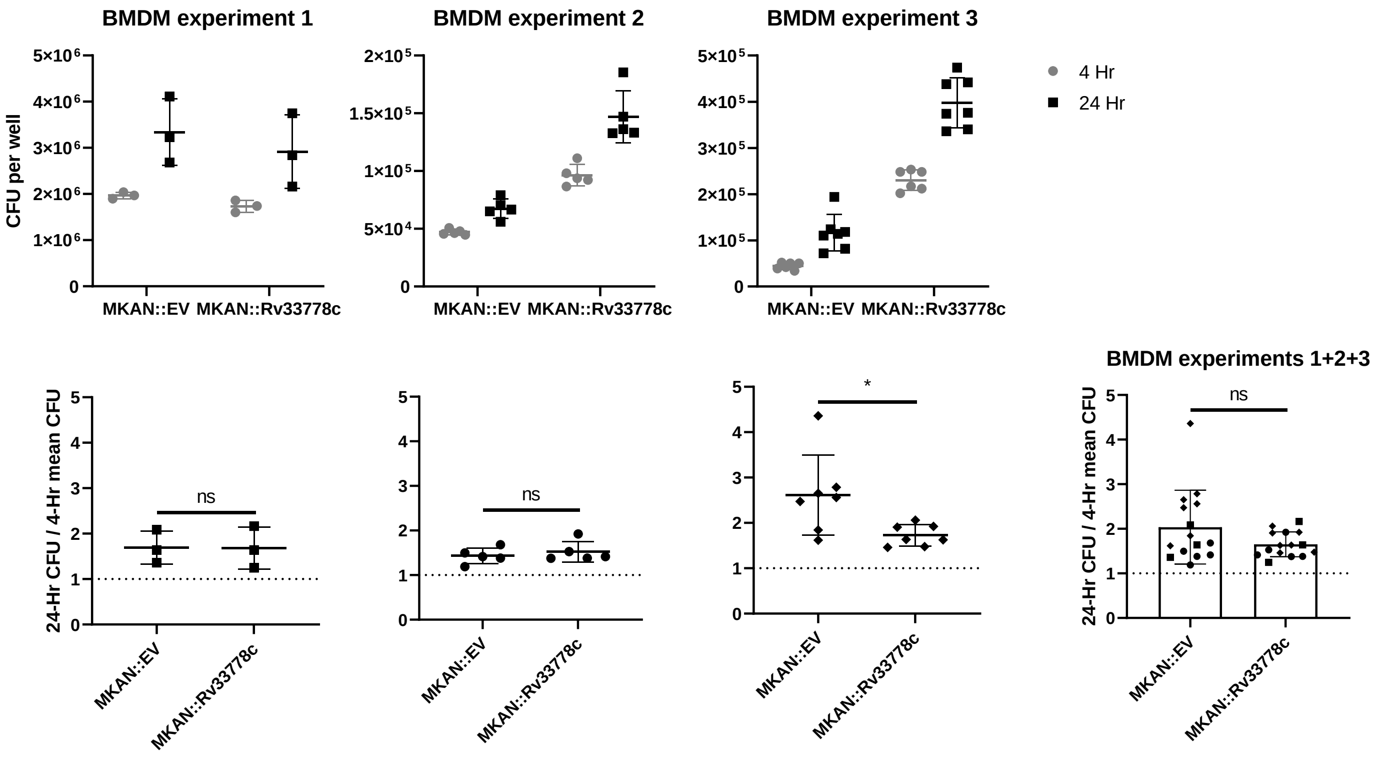

### Supplemental Figure 6

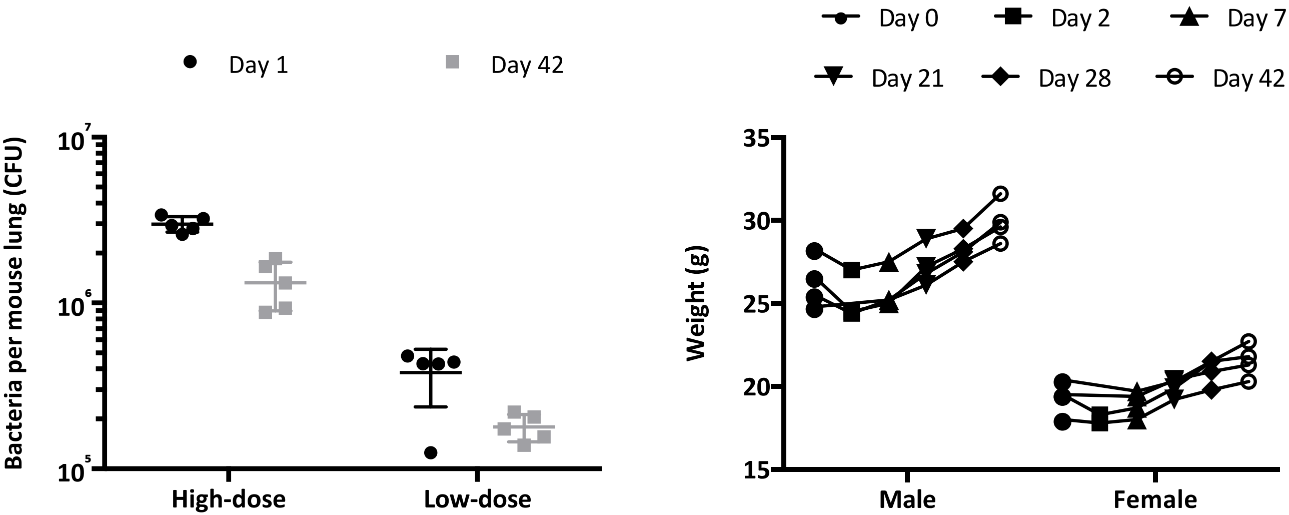

### Supplemental Figure 7

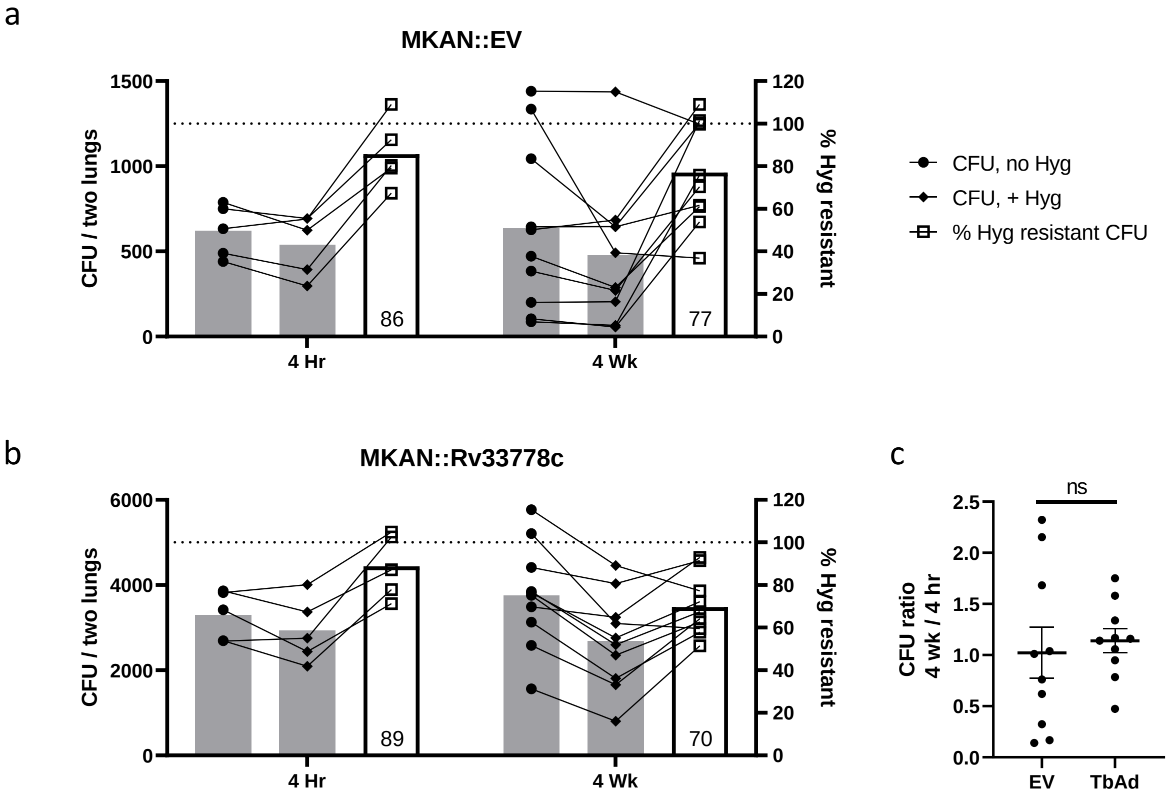
