## Supplemental Table 1 for "Heterologous production of 1-tuberculosinyladenosine in *Mycobacterium kansasii* models pathoevolution towards the transcellular lifestyle of *Mycobacterium tuberculosis*"

| Primer name | Primer sequence | Use |
|---|---|---|
| **BamHI-Rv3377-78c-F** | CGGGATCCATGAACTTGGTTAGCGAAAAAG | Cloning |
| **HindIII-Rv3377-78c-R** | CCAAGCTTTCATTGGTTACTCTCATCGACC | Cloning |
| **gDNA-Rv3377c-check-F** | TCGAGCACAGCCTATGACAC | Genotyping |
| **gDNA-Rv3377c-check-R** | ACCGATCCATTTGTCTCCTG | Genotyping |
| **gDNA-Rv3378c-check-F** | CACGAGGTCCACGTTCTTTT | Genotyping |
| **gDNA-Rv3378c-check-R** | CAACCCACACCGAAAACTCT | Genotyping |
| **Rv3377-78cSeq-F1** | GCCTTTGAGTGAGCTGATACC | Sanger sequencing |
| **Rv3377-78cSeq-F2** | GCTGGTTTCACCTCGAATG | Sanger sequencing |
| **Rv3377-78cSeq-F3** | GTCATTTCCGGCCCAAAC | Sanger sequencing |
| **Rv3377-78cSeq-F4** | CTTGGCGTACATCTCATCGA | Sanger sequencing |
| **Rv3377-78cSeq-R1** | GATAATCTCTCTCCGCGTG | Sanger sequencing |
| **Rv3377-78cSeq-R2** | CTTATTCGACGTGAGGCTG | Sanger sequencing |
| **Rv3377-78cSeq-R3** | CATCGTAGGCCACACTTGTG | Sanger sequencing |
| **Rv3377-78cSeq-R4** | CTGTCGTTCACGGCTCTAG | Sanger sequencing |
| **qMKANSigA-F** | CGGAGAAGGTGCTCGAAATC | qRT-PCR |
| **qMKANSigA-R** | TGGTCTGGTCCAGCGAGATC | qRT-PCR |
| **qRv3377c-F** | CAAGCTCTGGCGCATTGG | qRT-PCR |
| **qRv3377c-R** | GATCTGCGCCGACAAGGA | qRT-PCR |
| **qRv3378c-F** | CAACGATGCGGCTGAGTCT | qRT-PCR |
| **qRv3378c-R** | TTACCATGCGTTTCGTTCCA | qRT-PCR |
